# Laboratory Evolution Experiments Help Identify a Predominant Region of Constitutive Stable DNA Replication Initiation

**DOI:** 10.1101/757963

**Authors:** Reshma T Veetil, Nitish Malhotra, Akshara Dubey, Aswin Sai Narain Seshasayee

## Abstract

The bacterium *E. coli* can initiate replication in the absence of the replication initiator protein DnaA and / or the canonical origin of replication *oriC* in a *ΔrnhA* background. This phenomenon, which can be primed by R-loops, is called constitutive stable DNA replication (cSDR). Whether DNA replication during cSDR initiates in a stochastic manner through the length of the chromosome or at specific sites, and how *E. coli* can find adaptations to loss of fitness caused by cSDR remain inadequately answered. We use laboratory evolution experiments of *ΔrnhA-ΔdnaA* followed by deep sequencing to show that DNA replication preferentially initiates within a broad region located ∼0.4-0.7 Mb clockwise of *oriC.* This region includes many bisulfite-sensitive sites, which have been previously defined as R-loop forming regions; and includes a site containing sequence motifs that favour R-loop formation. Initiation from this region would result in head-on replication-transcription conflicts at rRNA loci. Inversions of these rRNA loci, which can partly resolve these conflicts, help the bacterium suppress the fitness defects of cSDR. These inversions partially restore the gene expression changes brought about by cSDR. The inversion however increases the possibility of conflicts at essential mRNA genes, which would utilise only a miniscule fraction of RNA polymerase molecules most of which transcribe rRNA genes. Whether subsequent adaptive strategies would attempt to resolve these conflicts remains an open question.

**Importance:** The bacterium *E. coli* can replicate its DNA even in the absence of the molecules that are required for canonical replication initiation. This often requires the formation of RNA-DNA hybrid structures, and is referred to as constitutive stable DNA replication (cSDR). Where on the chromosome does cSDR initiate? We answer this question using laboratory evolution experiments and genomics, and show that selection favours cSDR initiation predominantly at a region ∼0.6 Mb clockwise of *oriC.* Initiation from this site will result in more head on collisions of DNA polymerase with RNA polymerase operating on rRNA loci. The bacterium adapts to this problem by inverting a region of the genome including several rRNA loci such that head-on collisions between the two polymerases are minimised. Understanding such evolutionary strategies in the context of cSDR can provide insights into the potential causes of resistance against antibiotics that target initiation of DNA replication.

## Introduction

Canonical chromosome replication in the bacterium *Escherichia coli* is initiated by the specific recognition of repetitive short sequence motifs within the origin of replication *oriC* by the protein DnaA. This is followed by DNA unwinding and the synthesis of an RNA primer that can then be extended by the replicative DNA polymerase III (1). Replication proceeds bidirectionally outwards of *oriC* before terminating at a locus positioned diametrically opposite to *oriC* on the circular chromosome (2).

Bidirectional replication from a single *oriC* might have been the selective force behind the evolution of several organisational features of the genomes of bacteria, especially of those capable of rapid growth. These features include the encoding of highly expressed essential genes close to *oriC* to take advantage of the higher copy number of these loci while replication is in progress, and on the leading strand of replication to minimise the detrimental effects of head-on collisions between the DNA polymerase and RNA polymerases transcribing these genes (3). The positioning of such genes close to *oriC* is conserved, and more so in fast growing bacteria (4, 5). Repositioning of such genes away from *oriC* or on the lagging strand can be detrimental to fitness, especially in nutrient rich conditions (6–8).

Can the *oriC-*DnaA dependent mechanism of replication initiation in bacteria be dispensed with? Though DnaA is highly conserved across bacteria, it cannot be detected by sequence homology in a few (Supplementary table 1). Mitochondria are not known to use *oriC-*DnaA-based DNA replication initiation (9, 10). In *E. coli* the realisation that replication initiation by DnaA is sensitive to inhibition of translation resulted in the discovery of non-*oriC,* non-DnaA dependent “*<u>S</u>*table *<u>D</u>*NA *<u>R</u>*eplication” (SDR) (11).

Multiple broad types of SDR - each with its own set of genetic requirements - have been described. Inducible SDR (iSDR) requires the SOS DNA damage response (11, 12). Constitutive SDR (cSDR) is activated by processes that stabilise RNA-DNA hybrids or R-loops (11), such as the inactivation of (a) RnhA, the RNA-DNA hybrid nuclease RNaseHI (13); and that of (b) the topoisomerase I TopA, which results in hyper negative supercoiling and elevated occurrence of RNA-DNA hybrids (14) Excessive R-loops have also been proposed to occur in strains defective for Rho-dependent transcription termination (15–18), though to our knowledge Rho-dependent transcription termination has not been explicitly associated with cSDR. Inactivation of RecG, a helicase for RNA-DNA hybrids with roles in DNA recombination can also activate SDR (19–23). Very recently Raghunathan et al. demonstrated the role of the DNA methylase Dam in suppressing aberrant *oriC*-independent chromosomal replication, and showed that the deficiency of this protein conferred SDR, and that this mechanism is resistant to RNaseHI over-expression (24). We note here that DNA replication by SDR is under normal conditions sub-optimal relative to canonical DNA replication. At least one report has described nSDR, as a non-*oriC,* non-DnaA dependent mechanism of chromosome replication employed by *E. coli* cells transiently during the stationary phase (25).

In this paper, we focus on *ΔrnhA* induced cSDR in *ΔdnaA* mutants of *E. coli* K12. An important question in cSDR is: where does DNA replication initiate and what consequence does this have on chromosome organisation? The Kogoma group, employing traditional *<u>m</u>*arker *f*requency *<u>a</u>*nalysis (MFA), had identified five ‘*oriK’* loci at which replication might initiate (26). MFA analysis uses the argument that origin-proximal loci have a higher copy number than the rest of the chromosome in growing cells, even if they are not synchronised, to identify potential origins. Recently, Maduike et al. used deep sequencing based high resolution version of MFA to identify potential *oriK* sites, which were proximal to those identified by Kogoma’s group. The strongest signal in the Maduike et al. study mapped within the terminus of replication (27). Nishitani and colleagues cloned and screened for fragments of the *E. coli* chromosome with potential for autonomous self-replication, and thereby identified a cluster of fragments again from within the terminus (28). However, both Maduike et al. and Nishitani et al. appear to agree that the terminus sites identified in their studies are not bonafide *oriK* sites (27, 28). In the Maduike et al. study, these terminus signals disappeared in a *Δtus* background in which replication forks trapped within the terminus are released. The authors conclude that the terminus signal may represent trapping of forks originating from initiation sites elsewhere on the chromosome (27). Some of the *ter* sites identified by the Horiuchi group lost their activities in *Δtus*, but others did not. The Horiuchi group argued that increased copy number of fragments from the terminus can be attributed to homologous recombination based events and not autonomous replication (28). Gowrishankar has synthesised these arguments (29), and in conjunction with his lab’s finding that RNA-DNA hybrids can occur throughout the chromosome (30), presented the case that cSDR can initiate anywhere on the chromosome; individual cells can initiate replication at different sites thus generating population-level heterogeneity; and these can well explain the prominent MFA signal within the terminus. In a recent paper, Brochu et al. argue that *ΔtopA-topB* (more so than *ΔtopA-rnhA*) cSDR cells show a strong copy number peak within the terminus suggesting an *oriK* site here, but do not evaluate it in a *Δtus* background (31). These authors however observe that the *ter* peak is maintained in a strain with a large inversion around the *ter*, arguing against this peak being merely a consequence of replication fork trapping events.

Here we attempt to answer the question of the existence of preferred *oriK* sites by taking the position that peak identification in high resolution MFA studies of cSDR is complicated by the slow growth phenotype of the parent strain, which results in weak origin to terminus copy number gradients. We address this using laboratory evolution experiments, generating suppressors that can generate strong copy number gradients even under the cSDR regime, while also identifying a principle underlying the suppression of the slow growth phenotype of cSDR.

## Results

### Next generation sequencing based MFA of *ΔrnhA-ΔdnaA* strain of *E. coli* K12

The gene *rnhA* encodes the RNaseHI nuclease that removes RNA-DNA hybrids. The *ΔrnhA* mutant displays cSDR and therefore suppresses the lethality of *ΔdnaA* and *ΔoriC* mutants (13). We obtained a *ΔrnhA* single deletion mutant by homologous recombination, and a *ΔrnhA-ΔdnaA-pHYD2388* (*dnaA*+*lacZ*+) mutant of *E. coli* K12 (MG1655) from Prof. J. Gowrishankar’s lab (24). To obtain *ΔrnhA-ΔdnaA*, we plated overnight cultures of *ΔrnhA-ΔdnaA-*pHYD2388 (*dnaA*+*lacZ*+) on X-gal agar plates. Spontaneous loss of the *dnaA*+ pHYD2388 plasmid produced white colonies (*dnaA*-*lacZ*-), which we selected and propagated as the *ΔrnhA-ΔdnaA* strain. *ΔrnhA* and *ΔrnhA-ΔdnaA* showed growth characteristics consistent with prior literature (Supplementary Figure 1).

In the rest of this manuscript, we use ‘*ori’* as an umbrella term, when required, to refer to all sites at which replication initiates: this may include *oriC* itself or *oriK* sites at which cSDR initiates. The terminus is a more complex sequence with multiple, directional replication termination motifs at which the Tus protein traps moving replication forks; we use the generic term ‘*ter’* to refer to the locus bounded by these termination motifs.

We isolated genomic DNA from *rnhA*+*dnaA*+, *ΔrnhA* and *ΔrnhA-ΔdnaA* strains of *E. coli* grown to exponential phase - corresponding to the culture’s highest growth rate - in LB. We sequenced the DNA libraries prepared from these samples to an average coverage of ∼200x on the Illumina platform. As controls, we sequenced DNA isolated from stationary phase populations. For *rnhA*+*dnaA*+, we observed a copy number gradient decreasing from *oriC* towards *ter*, symmetrically on either side of *oriC*, such that the number of reads mapping around *oriC* was 2.48 fold higher than that around *ter* (Figure 1A). The corresponding plot for stationary phase cells was relatively flat (Figure 1A lower panel).

**Figure 1:**
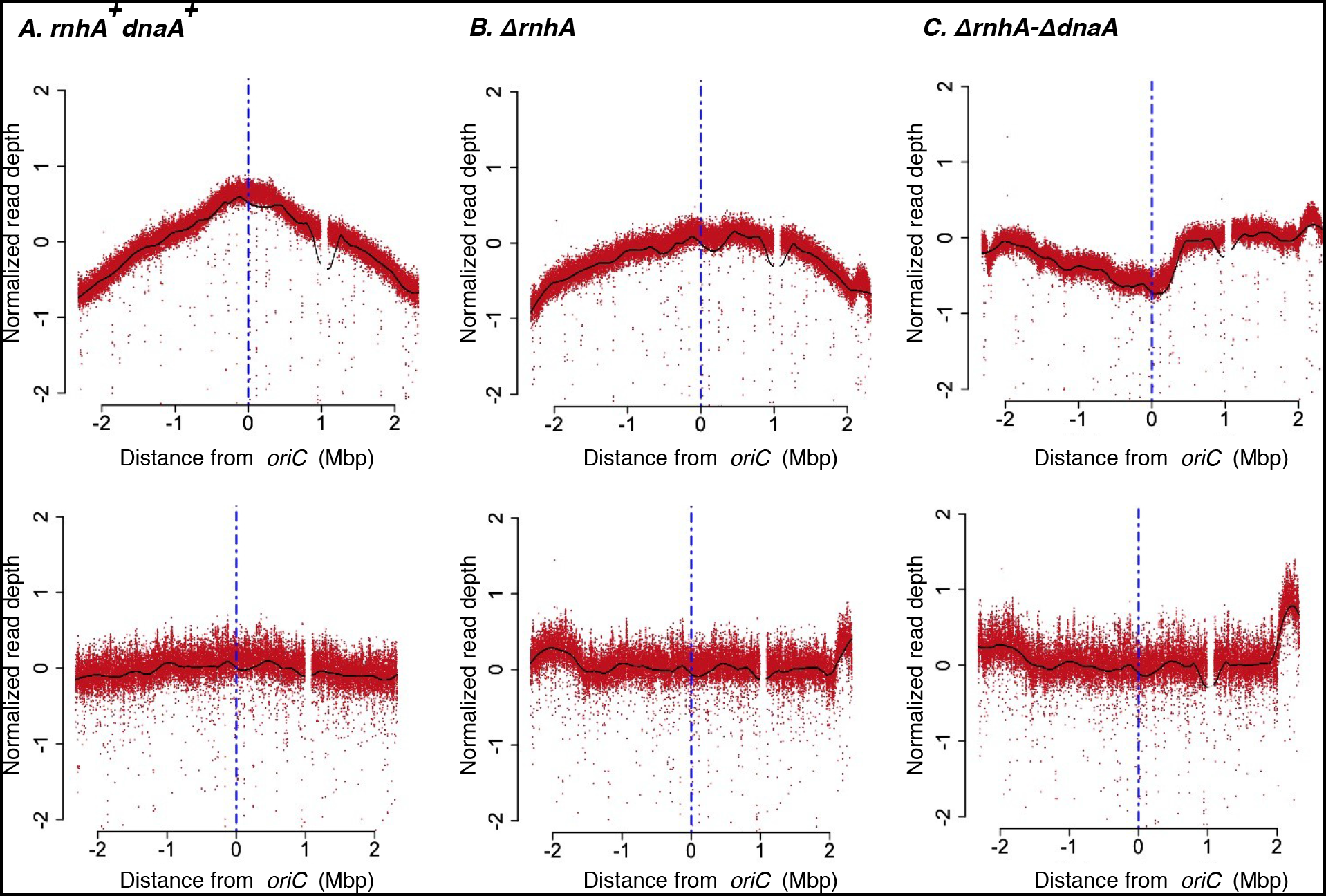
Deep sequencing based MFA plots for *rnhA^+^dnaA^+^, ΔrnhA and ΔrnhA-ΔdnaA*. The upper panels show the MFA plots for (A) *rnhA^+^dnaA^+^,* (B) *ΔrnhA* and (C) *ΔrnhA-ΔdnaA* at the exponential phase of growth and the lower panels show the same for the stationary phase. The X-axis represents the distance of a locus either side of *oriC* (in Mbp), with *oriC* itself being the centre (blue vertical line). The Y-axis represents the log_2_ values of frequency of reads divided by the mode of the distribution of read counts (see methods).

The *ΔrnhA* mutant, in which both *oriC-*DnaA-dependent replication initiation and cSDR are active also showed a fairly steep gradient (Figure 1B). In line with its slow growth, *ΔrnhA-ΔdnaA* showed a flat curve with a few peaks which are candidates for *oriK* sites (Figure 1C). The strongest peaks were those present within the t*er*, which has been rejected previously (27, 28) as arising as a consequence of trapping of replication forks initiating elsewhere on the chromosome; and a second site around 0.5-0.6Mb clockwise of *oriC*, which we call as *oriK45* for it being located at approximately 4.5 Mb into the genome sequence of *E. coli K12 MG1655.* In large part, the patterns observed were consistent with those observed by Maduike et al. (27) and Dimude et al. (23) (Figure 1C and Table 1).

**Table 1:** All numbers are genome coordinates (bp). The *oriC* peaks, when present, are not included in this table. Genomic coordinates mentioned for Maduike et al., 2014 is normalized to the respective positions of peaks in *E.coli* K12 MG1655 genome version NC_000913.3.

| Strain | Identified Peaks |  |  |  |  |  |  |
| --- | --- | --- | --- | --- | --- | --- | --- |
| $\Delta rnhA\text{-}\Delta dnaA$ | 531400 | 1449000 | 1969800 | 2988600 | - | 3699200 | 4546800 |
| $\Delta rnhA\text{-}\Delta dnaA$<br>(Maduike et al. 2014) | 790277 | 1481276 | 1869776 | - | 3228978 | - | 4538577 |
| $\Delta rnhA$ | - | 1453600 | - | - | - | - | 4379400 |
| $\Delta rnhA$<br>(Maduike et al. 2014) | - | 1471624 | - | - | - | - | 4253823 |

Whether *oriK45* is a genuine replication initiation site, whether it is indeed a ‘preferred’ site, and whether other minor peaks around the chromosome can represent substantial *oriK*s remain complicated to answer with the present dataset. This is at least in part because of the slow growth phenotype of the mutant which ensures that there is hardly any *ori*-*ter* copy number gradient even during periods of its highest growth rate.

### Laboratory evolution experiments of *ΔrnhA-ΔdnaA*

To obtain cSDR strains that grow fast and therefore display strong *ori*-*ter* gradients, we performed laboratory evolution experiments in which *ΔrnhA-ΔdnaA* was iteratively diluted into fresh LB and grown to saturation. We used eight independent lines, each derived from a single *ΔrnhA-ΔdnaA* colony to 36 rounds of dilution and growth, corresponding to an estimated 288 generations. Over time, the growth of the population substantially improved (Figure 2).

**Figure 2:**
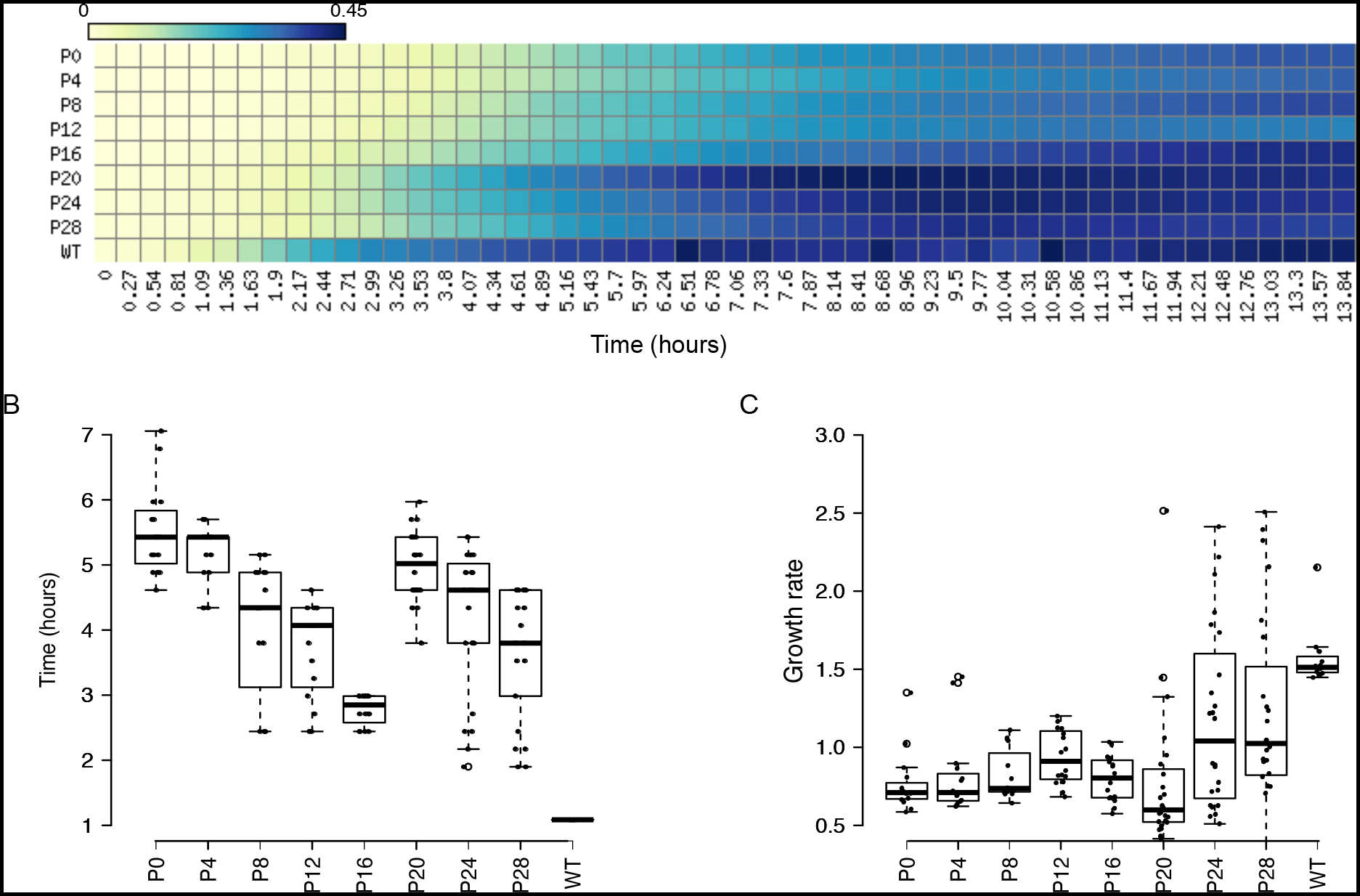
Growth characteristics of evolved mutants: (A) Heat map representing growth of an independently evolved population of *ΔrnhA-ΔdnaA* from passage 0 (P0) to passage 28 (P28) based on OD measurements. X-axis shows time in hours, Y-axis shows the number of passage and the colours represent mean OD values as indicated in the colour bar. Similar growth characteristics were observed for other evolution lines. (B) and (C) box plots for lag time and growth rate followed by all independently populations respectively. Passage 28 population shows a significantly greater growth rate than that of parental (P0) strains (P<<< 0.001, Wilcoxon test, one tailed).

We plated aliquots of the culture after each day and noticed the presence of colonies that were visibly larger than those of the parent *ΔrnhA-ΔdnaA*. We randomly picked 54 colonies of varying sizes - sampling across 3 independently evolved populations and 5 time-points and including the zeroth day populations - and subjected their genomic DNA to Illumina sequencing. Similar to our sequencing runs with the parent *ΔrnhA-ΔdnaA*, we sequenced DNA isolated from mid-exponential phase. Stationary phase DNA sequencing was performed for a select few colonies based on genotypes identified from exponential phase DNA sequencing.

For all these strains, we calculated the ratio between the maxima and the minima of the mid-exponential phase copy number graphs (see Materials and Methods), and found that this ratio ranged between 0.86 and 2.8 (Supplementary table 2). At the lower end, a few colonies showed gradients not too different from the *ΔrnhA-ΔdnaA* parent. The steepest gradients approached, but rarely matched that of *rnhA*+*dnaA*+.

**Table 2:** Probability of head-on-collisions were predicted for 3.3Mbp to 4.25Mbp region of the chromosome which includes the inverted region of *ΔrnhA-ΔdnaAinv*^rrnD-rrnE^ strain. The values were calculated by taking the ratio of sum of RNA coverage values of all genes on the lagging strand with respect to the single predominant *ori* position to the total RNA coverage for the region(P(HO) = sum(lagging strand RNA coverage)/ total RNA coverage). The analysis was done for different classes of genes separately by using the functional annotations for genes from NC_000913.3(.ptt) file.

| Strain | Head-on-collision rate |
| --- | --- |
| **All genes** |  |
| *rnhA<sup>+</sup>dnaA<sup>+</sup>* | 0.31 |
| *ΔrnhA-ΔdnaA* | 0.67 |
| *ΔrnhA-ΔdnaA<sup>rrnD-rrnE</sup>* | 0.39 |
| **ribosomal genes** |  |
| *rnhA<sup>+</sup>dnaA<sup>+</sup>* | 0.22 |
| *ΔrnhA-ΔdnaA* | 0.84 |
| *ΔrnhA-ΔdnaA<sup>rrnD-rrnE</sup>* | 0.24 |

### Large inversions around *oriC* suppress the growth defect of *ΔrnhA-ΔdnaA*

We next used these sequencing data to identify mutations – both point variations including indels, as well as structural variations such as large amplifications, deletions and inversions. Large amplifications and deletions can be identified by sharp local increases or decreases respectively in copy number. Inversions can be detected as local flips in copy number plots of exponential phase genomic DNA sequencing data with clear *ori*-*ter* gradients (32). We found several point mutations in the evolved clones, not present in the *ΔrnhA-ΔdnaA* parent (Figure 3 and Supplementary figure 2). ∼90% of colonies carried a mutation upstream of one of two rRNA operons, *rrnD* and *rrnC.* One clone carried an in-frame deletion mutation in *tus (*Δ6 bp (1,684,458-1,684,463), which translates to a QSL-L variation. We did not find any amplification, and the only deletion that was apparent in the data was an ∼97 kb ([*mmuP*]–[*mhpD*]) deletion around the *lac* locus, which is part of the genotype of the *rnhA*+*dnaA*+ founder strain used in this study (24).

**Figure 3:**
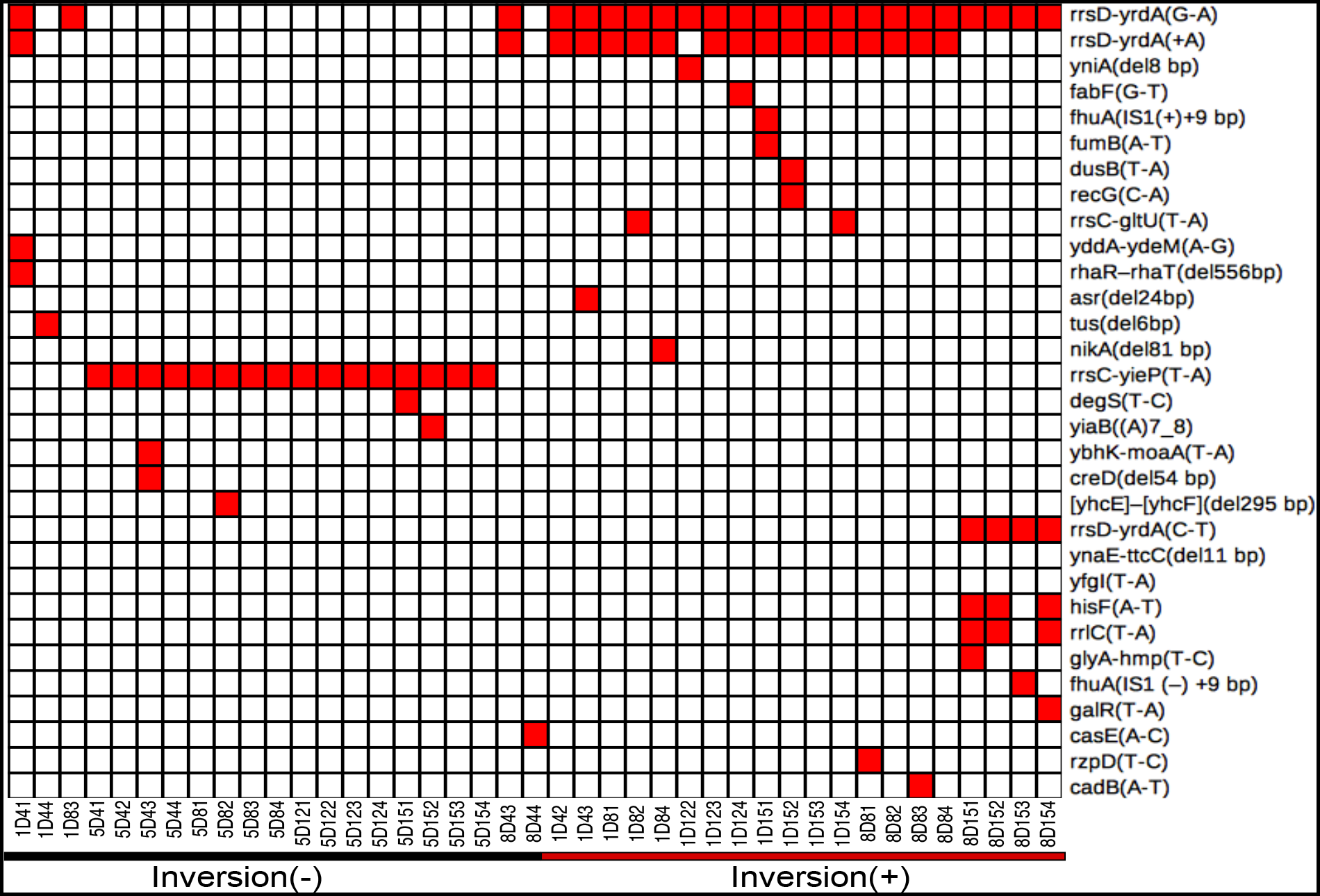
Unique mutations in suppressor mutants. Heat map representing unique mutations (100% frequency) in all independent colonies sequenced generated using matrix2png. Colour represents the presence of a mutation in the respective gene shown on Y-axis. X-axis represents sample IDs of suppressor mutants evolved from three independent populations. Presence and absence of chromosomal inversions are represented using a red and black lines respectively. Mutations in pseudogenes and those also found in parental strains are listed in Supplementary Figure 2.

We found inversions around *oriC* in ∼45% of the evolved colonies (Figure 3 and Figure 4). One end of these inversion was *rrnD*, located 3.42 Mb counter-clockwise of *oriC* in the reference genome of *E. coli* K12 MG1655. In ∼80% of inversions, the other end was *rrnC* (3.94 Mb), and in the remaining, the second end was *rrnE* (4.2Mb). The *rrnD-rrnC* inversion (*ΔrnhA-ΔdnaA-inv*rrnD-rrnC) measured ∼0.5 Mb and the *rrnD-rrnE* (*ΔrnhA-ΔdnaA-inv*rrnD-rrnE), ∼0.8 Mb (Figure 4). We used long read nanopore sequencing to assemble the genome of the clone with the longer *rrnD-rrnE* inversion into just one contig *de novo*, and confirmed the presence of the inversion (Figure 5A).

**Figure 4:**
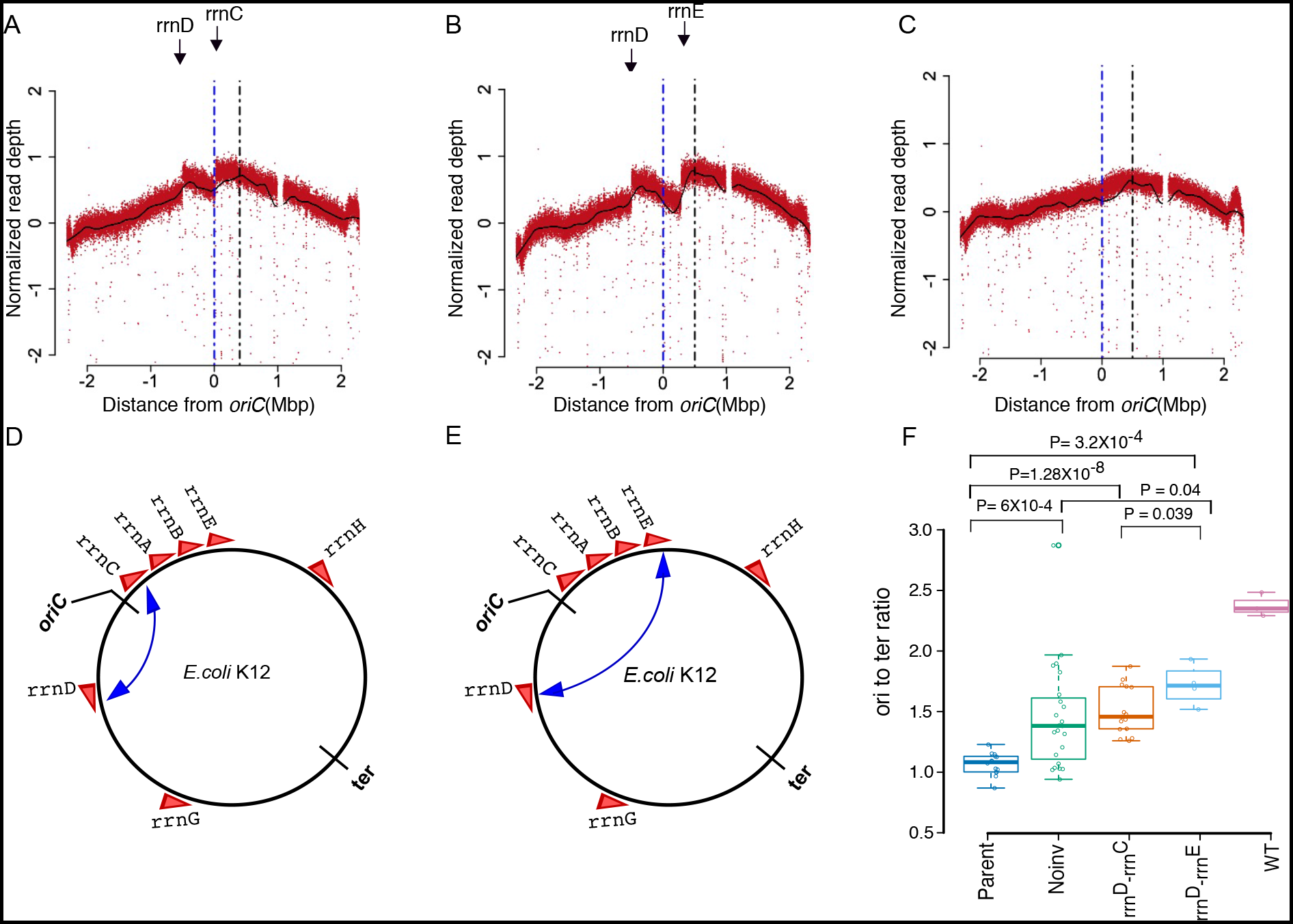
Deep Sequencing based MFA plots for suppressor mutants. (A), (B) and (C) represents MFA plots for (A) *ΔrnhA-ΔdnaAinv^rrnD-rrnC^*, (B) *ΔrnhA-ΔdnaAinv*^rrnD-rrnE^, (C) *ΔrnhA-ΔdnaANoinv* sequenced at the exponential phase of growth. The dotted blue line represents the *oriC* position and the black line represents the position at maxima of Loess fit value. (A) and (B) plots show the presence of different chromosomal inversions flanked by *rrn* operons (mentioned above) and the position of inversion on the chromosome has been schematically represented here(D and E). (F) box plot representing *ori-to-ter* ratio differences in different populations of evolved clones compared to wild type *E.coli*. (X-axis labels; Parent-*ΔrnhA-ΔdnaA* strain passage 0 clones, No inv: suppressor mutants which do not show the presence of chromosomal inversion, rrnD-rrnC: Clones which shows presence of a chromosomal inversion from rrnD-rrnC (*ΔrnhA-ΔdnaAinv^rrnD-rrnC^*) and rrnD-rrnE: Clones which shows presence of a chromosomal inversion from rrnD-rrnE(*ΔrnhA-ΔdnaAinv^rrnD-rrnE^*).

**Figure 5:**
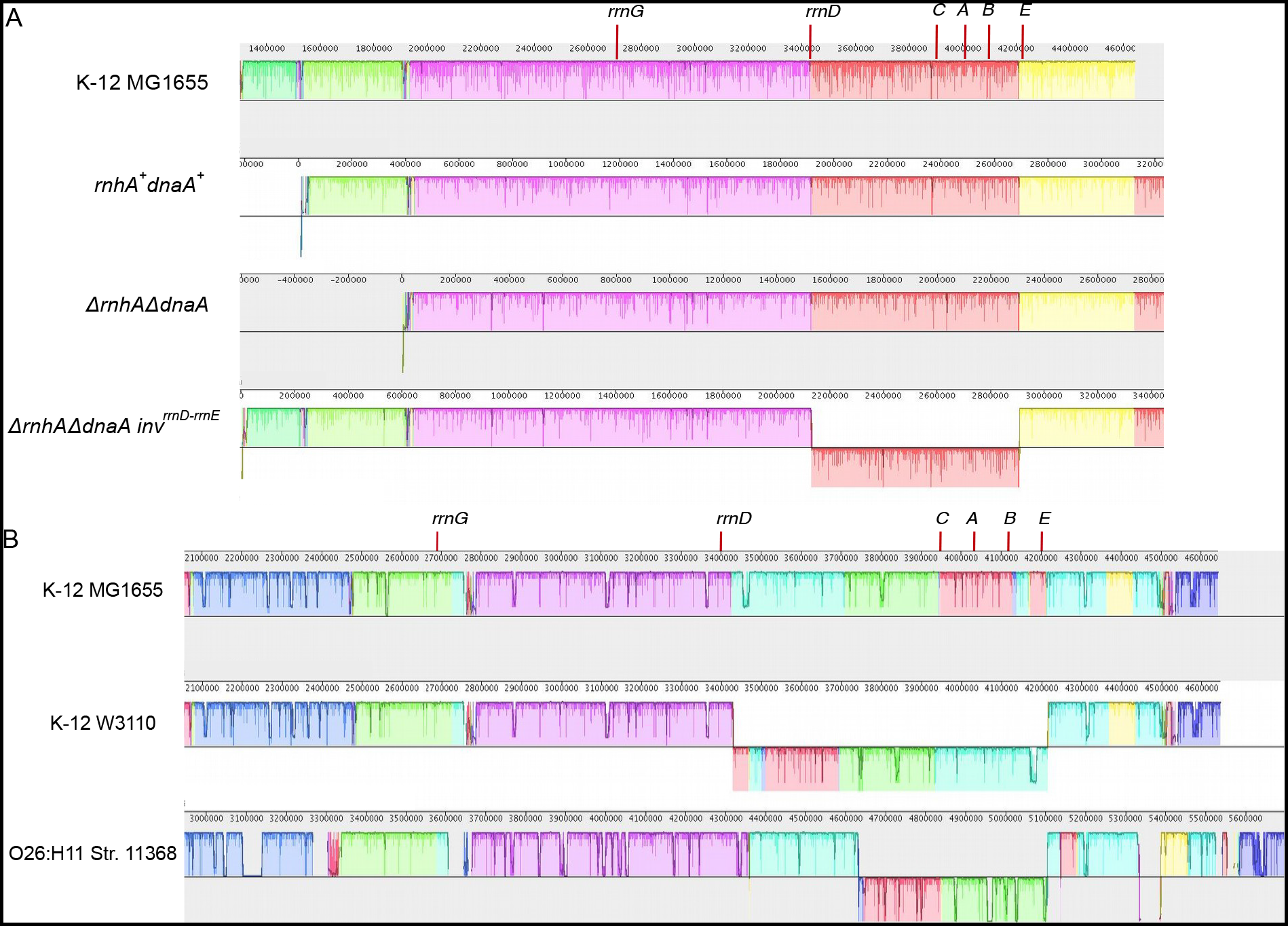
(A) Plot showing sequence alignment of denovo assembled contigs obtained from Nanopore sequencing for *rnhA^+^dnaA^+^, ΔrnhA-ΔdnaA* and *ΔrnhA-ΔdnaAinv^rrnD-rrnE^* strains with respect to *E.coli* K12 MG1655 genome. Strand shift of red color bar for the *ΔrnhA-ΔdnaAinv^rrnD-rrnE^* shows the presence of ∼0.8 mb chromosomal inversion. Position of *rrn* operons near *ori* region is marked. (B) Figure representing sequence alignment (using Mauve) of selected region of chromosome for three *E.coli* strains. *E.coli* K12 W3110 and *E.coli* O26:H11 str.11368 shows the presence of a chromosomal inversion around *oriC* region compared to the reference *E.coli* K12 MG1655. Strand shift for the colored bars represents chromosomal inversion. Position of *rrn* operons near *ori* region is marked.

Thus both inversions would move a set of rRNA operons from clockwise to counter-clockwise of *oriC*, and the *rrnD* operon in the opposite direction. Irrespective of the presence of the inversion, all these rRNA operons would continue to lie on the leading strand of canonical replication from *oriC*. That the fitness cost of these inversions would be minimal under conditions of normal DNA replication is also suggested by the fact that inversions bounded by at least one *oriC*-proximal rRNA operon are found in 37 other *E. coli* genomes (out of 675 considered), including another strain of *E. coli* K12 (W3110) (33) (Figure 5B and Supplementary table 3). Colonies with either inversion in the present study also carried the following mutations upstream of *rrnD*: (a) G-A (position 3,429,052) and +A (3,429,054) or (b) C-T (3,429,055) (Figure 3).

We then compared the *maximum-minimum* ratios in the copy number plots of clones (not considering the peak within *ter*) with the two types of inversions and those without. For this analysis, we grouped all colonies without an inversion together, fully aware that this is a genetically heterogeneous group. Clones with the longer *rrnE-rrnD* inversion showed significantly higher maximum-minimum ratios than those with the shorter *rrnC-rrnD* inversion (P = 0.039, Wilcoxon test one-tailed) (Figure 4F). Therefore, the longer inversion appears to be a better suppressor of the growth defect of cSDR than the shorter inversion. Many clones without the inversions, including the one with the Δ6 bp inframe deletion mutation in Tus, showed substantially smaller maximum-minimum ratios, though a few colonies did show higher values.

### *oriK45* as a preferred initiation site for cSDR in suppressors

We identified the locations of the maxima of the copy number curve for the suppressors, while ignoring the *ter* peak. We noticed that these, across all cSDR strains used in this study, mapped to ∼4.3 Mb to 4.6 Mb clockwise of *oriC*, in proximity to *oriK45* (Figure 6A). Consistent with this, all suppressors showed a copy number peak at *oriK45* (Figure 6A and Supplementary table 4). This suggests that *oriK45* is a predominant site of cSDR initiation in *all* suppressors identified here.

**Figure 6:**
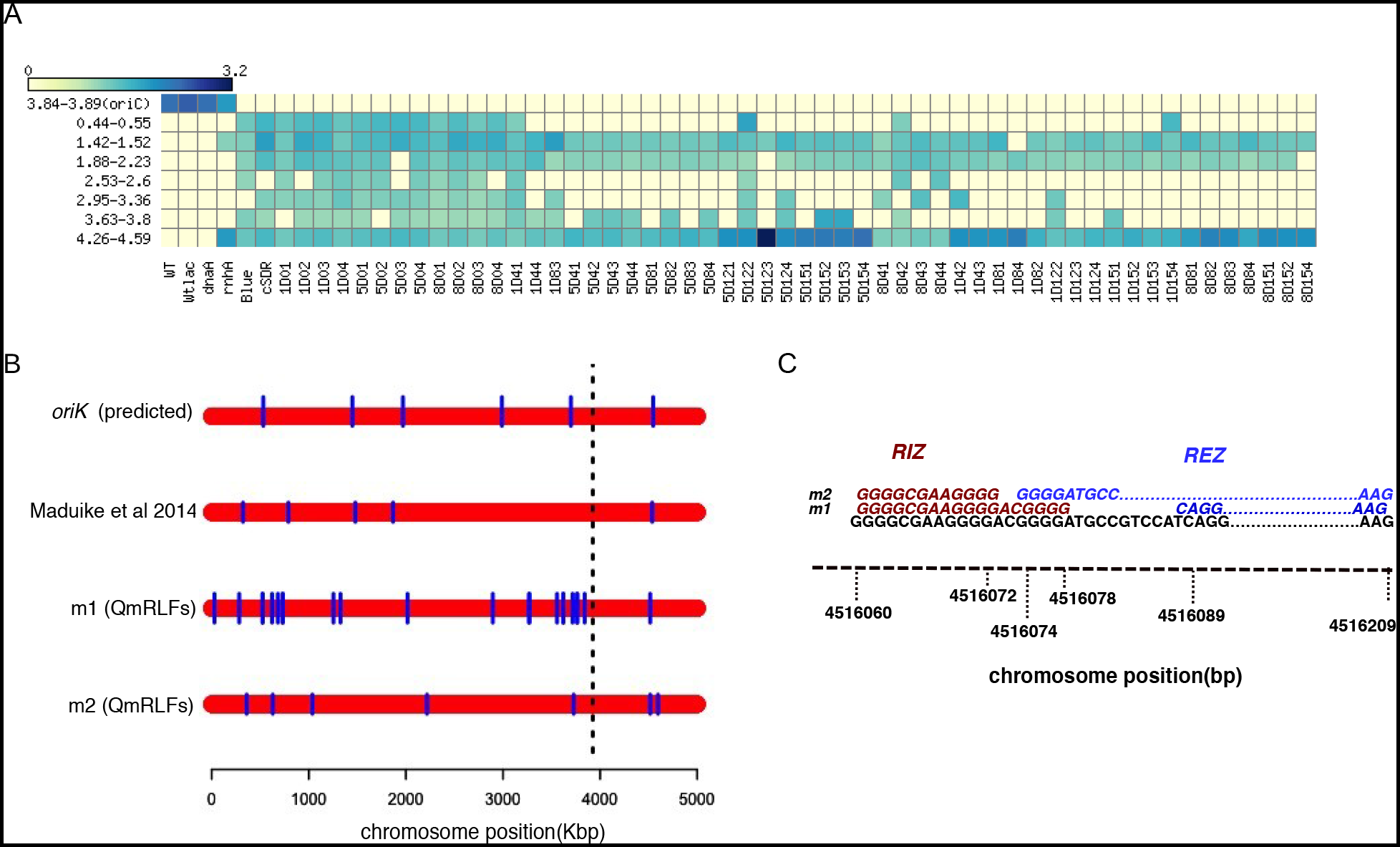
*oriK45* as a preferred initiation site for cSDR in suppressor mutants: (A) heatmap showing predicted *oriK* position ranges from marker frequency analysis across evolved strains. Y-axis represents the chromosomal positions of predicted *oriK* ranges in Mbp. Colour indicates the ratio of corresponding LOESS smoothed normalised read count to the LOESS minima of the plot at the peak. (B) plot showing positions of R-loops predicted by m1 and m2 model of QmRLFs on *E.coli* chromosome in comparison with position of predicted *oriK* sites in *ΔrnhA-ΔdnaA* strain from Maduike et al. 2014 and the *oriK* sites for the same strain mentioned in this study. Each red bar represents the bacterial chromosome on which the R-loops positions are marked in blue lines. (C) Represents the sequence motif of R-loop initiation zone (RIZ) and R-loop elongation zone predicted by m1 and m2 models of QmRLFs around 4.51mb region of *E.coli* K12 MG1655 chromosome.

In the strongest suppressors, we observed a strong copy number gradient peaking at *oriK45* and declining towards *ter* (figure 4A and 4B). (DNA copy number curves for all evolved mutants are available at https://doi.org/10.6084/m9.figshare.11800299) The peak in *ter* was computationally detected in all suppressors. However, this peak was weak in two of the suppressors. One of these contained a 6 bp in-frame deletion mutation in *tus* (figure 3, sample ID 1D4_4), and displayed a copy number pattern similar to that observed for *Δtus* by Maduike et al., (27) indicating that the mutation observed here causes loss of function. This strain did show a slight copy number peak at *oriK45,* but being a relatively weak suppressor does not permit a more confident assignment.

### *oriK45* is proximal to predicted R-loop-forming sites

We asked whether *oriK45* is proximal to regions with high propensities to form RNA-DNA hybrids. We used a computational technique that searches for two G-rich patterns on a given DNA sequence to identify loci that have the propensity to form RNA-DNA hybrids (34, 35). This method predicted ∼30 R-loop favouring sites, showing homology to at least one of the two RNA-DNA hybrid-forming sequence patterns, across the *E. coli* chromosome (Figure 6B). 8 of the 11 copy number bumps described by us or by Maduike et al. for *ΔrnhA-ΔdnaA* were within 200 kb of at least one of the predicted sites. This is statistically significant compared to random assignment of genome coordinates to experimentally predicted copy number peaks (*P =* 10-5, Z-score, permutation test across 1,000 repetitions, one-tailed). However, only one site showed homology to both RNA-DNA hybrid-forming sequence patterns; this site is at 4.51 Mb (Figure 6C), within the range defined by *oriK45*.

Krishna Leela et al. (30) had identified bisulfite sensitive regions of *E.coli* chromosome and defined these as preformed R-loops. However, we did not find any statistically significant overlap of these sites with *oriK45*. Nevertheless, we found two clusters of highly bisulfite-sensitive genes in the *oriK45* region, and also observed that the R-loop forming sequence mentioned above was also highly bisulfite sensitive.

Nishitani et al., while screening for genomic DNA fragments capable of autonomous replication, describe a site called *hotH*, which is at 4.55-4.56 Mb (28). However, to our knowledge, these authors did not report further exploration of the *hotH* site and focussed instead on the characterisation of the cluster of fragments from within *ter*. Among the transposon insertions found to affect replication of *ΔtopA*-mediated cSDR is an insertion within *fimD*, which is again in the region defined by *oriK45* (36).

### cSDR from *oriK45* has pleiotropic effect on gene expression

What are the effects of cSDR on gene expression - as measured by global patterns along the length of the chromosome, and signatures on pathways related to DNA replication, repair and transcription? To what extent does the suppression of growth defects of cSDR by the inversion around *oriC* reverse these effects? Towards answering these questions, we performed exponential phase transcriptome analysis of *rnhA*+*dnaA*+, *ΔrnhA, ΔrnhA-ΔdnaA, ΔrnhA-ΔdnaA-inv*rrnD-rrnC*, ΔrnhA-ΔdnaA-inv*rrnD-rrnE using RNA-seq.

Both *ΔrnhA* and *ΔrnhA-ΔdnaA* induced large changes in gene expression when compared to *rnhA*+*dnaA*+. 600 genes were up-regulated and 543 down-regulated by a log (base 2) fold change of 1.5 or above in *ΔrnhA-ΔdnaA.* The corresponding numbers for *ΔrnhA* are 472 and 360. Nearly 75% of all genes induced in *ΔrnhA* were also induced in *ΔrnhA-ΔdnaA*; the proportion for down-regulated genes being ∼80%. Despite the overlap in these gene lists, the magnitude of differential expression was in general less in Δ*rnhA* than in *ΔrnhA-ΔdnaA* (*P* < 10-10, paired Wilcoxon test comparing magnitudes of differential expression). Functional classification of differentially expressed genes using Clusters of Orthologous Groups (COGs) showed an enriched up-regulation (*P <* 0.01, Fisher’s Exact test) of various classes of genes including replication recombination and repair genes, ion transport and metabolic genes and translation genes (Supplementary table 5). On the other hand, cell motility, energy production and conversion, carbohydrate transport and metabolic genes showed a significant down-regulation in *ΔrnhA-ΔdnaA*.

Genes encoding several members of the SOS response, including the cell division inhibitor SulA, error prone polymerases DinB and UmuC, RuvB and C are up-regulated in both *ΔrnhA* and *ΔrnhA-ΔdnaA*. *dinF,* the SOS inducible gene that also confers protection against oxidative stress was induced in both the mutants. Other signatures for an oxidative stress response included the induction *of sufB-E*, whose protein products are involved in iron-sulfur cluster biogenesis under oxidative stress (37). Very few members of the general stress response (∼6%; under-represented when compared to Sigma70 targets, *P* = 4 x 10-6, Fisher’s Exact Test), defined as targets of Sigma38 (RpoS), were induced.

We also observe an up-regulation of *holB* and *holD*, encoding the delta-prime and the epsilon subunits respectively of the replicative DNA polymerase III. This might in part be consistent with the SOS response, in light of the evidence that induction of SOS responsive DNA polymerases can be lethal in a genetic background that is defective for HolD (38). The gene *topA*, encoding topoisomerase, which can decrease R-loop formation presumably through its DNA relaxing activity, is also up-regulated.

We observe that several genes encoding components of the ribosome are up-regulated in the inversion mutants. At least three DEAD box RNA helicase genes (*rhlE*, *dbpA* and *srmB*) that are involved in ribosome assembly are also up-regulated. Finally, *rapA*, the gene encoding the RNA polymerase recycling factor ATPase, which is required for reloading stalled RNA polymerase is up-regulated.

### Gene expression changes show limited but significant correlation to DNA copy number changes

Overall, there is a gradient - decreasing from *oriC* towards *ter* - in the fold change in gene expression between *rnhA*+*dnaA*+ *and ΔrnhA-ΔdnaA*. In other words, genes that are proximal to *oriC* (and *oriK45*) are more strongly down-regulated in *ΔrnhA-ΔdnaA* when compared to *rnhA*+*dnaA*+ (Supplementary figure 3). At this level, the fold change in *rnhA*+*dnaA*+, in relation to *ΔrnhA-ΔdnaA*, shows strong similarity to that in *ΔrnhA* and *ΔrnhA-ΔdnaA-inv*rrnD-rrnE (Pearson correlation coefficient = 0.64 for both comparisons), and slightly less similar to *ΔrnhA-ΔdnaA-inv*rrnD-rrnC(Pearson correlation coefficient = 0.55) (Figure 7). These indicate that a portion of the gene expression change in *ΔrnhA-ΔdnaA* relative to *rnhA*+*dnaA*+ is reversed by the longer inversion *ΔrnhA-ΔdnaA-inv*rrnD-rrnE, and probably less so by the shorter inversion *ΔrnhA-ΔdnaA-inv*rrnD-rrnC. Nevertheless, the magnitude of the difference in gene expression between *rnhA*+*dnaA*+ and *ΔrnhA-ΔdnaA* is higher than that between the suppressors and *ΔrnhA-ΔdnaA* (*P* < 10-10, paired Wilcoxon test comparing magnitudes of differential expression).

**Figure 7:**
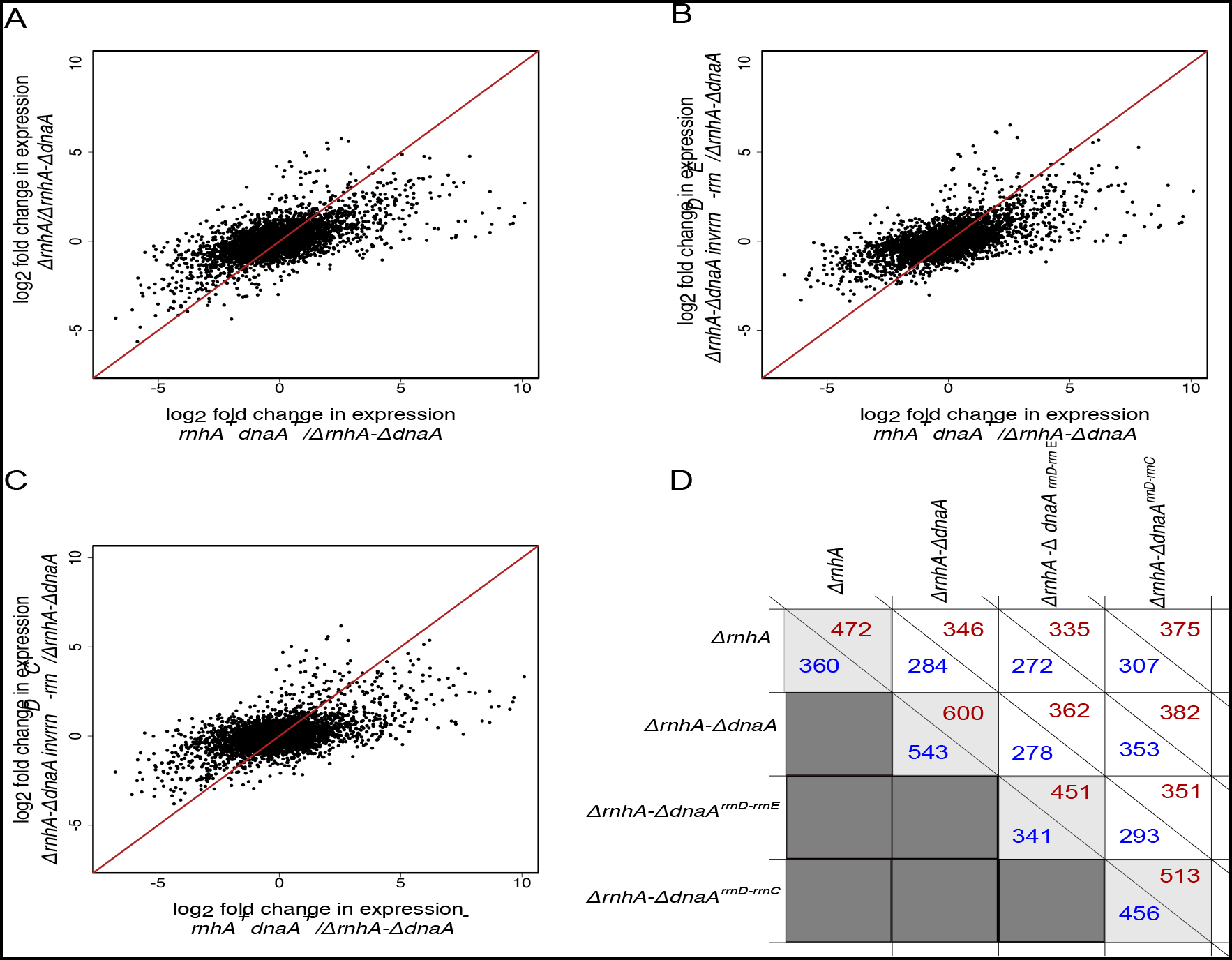
Scatterplots representing correlation of log2 fold change in gene expression for different conditions, compared to *ΔrnhA-ΔdnaA* strain. (A) *ΔrnhA* vs *rnhA^+^dnaA^+^* (B) *ΔrnhA-ΔdnaAinv^rrnD-rrnE^* vs *rnhA^+^dnaA^+^* and (C) *ΔrnhA-ΔdnaAinv^rrnC-rrnE^* vs *rnhA^+^dnaA*^+^. The pearson correlation values for (A), (B), (C) are 0.638, 0.639, and 0.553 respectively. (D) plot representing the number of up-regulated(red) and down-regulated(blue) genes for all strains compared with *rnhA^+^dnaA^+^*.

A small, but statistically significant portion of the difference in gene expression can be explained by differences in DNA copy number – a consequence of differences in maximal growth rates – as measured by NGS sequencing of matched exponential phase genomic DNA samples (Pearson correlation coefficient ∼ 0.2, *P* < 10-10). These correlations between DNA copy number and RNA-seq based gene expression fold changes increase to over 0.75 in all comparisons when gene expression data are smoothed by LOESS, which averages out local variation in expression levels.

The movement of the origin of replication to *oriK45,* and the large inversion, might affect the macrodomain structure of the chromosome (39), as well as supercoil gradients (40). *oriK45* would be located at the right extreme of the *o*ri macrodomain. The left end of the larger inversion is within a non-structured region of the chromosome, whereas the right end is within the *ori* macrodomain, and such an inversion could have consequences to cell physiology as well as gene expression (41).

What the precise effect of these chromosome structure parameters are on the transcriptional profile is, in the absence of chromosome conformation data under cSDR, is not clear at the moment.

Therefore, overall gene expression changes along the chromosome are weakly correlated with the distance of a gene from *oriC* (and *oriK45*) and changes in DNA copy number. Gene expression changes that occur in *ΔrnhA-ΔdnaA* relative to *rnhA*+*dnaA*+ are partly compensated by inversion containing suppressors.

### The inversions reduce replication-transcription conflicts at rRNA loci but not at essential mRNA genes

To understand the impact of inversions on transcription-replication collisions, we calculated a fractional score for the occurrence of head-on collisions for genes on the lagging strand with respect to replication from *oriC* or *oriK45* using RNA sequencing data (see materials and methods). This score was lowest at 0.31 for *rnhA*+*dnaA*+. This increased to 0.67 in *ΔrnhA-ΔdnaA*, but was reduced to 0.39 in the suppressor *ΔrnhA-ΔdnaAinv*rrnD-rrnE (Table 2). This effect was the strongest when only rRNA genes (5S rRNA, which is not depleted as part of the RNA prep experiment) were considered. Despite the large decrease in replication-transcription conflict in the inversion-containing suppressors, the activation of the SOS response in cSDR is not reversed, even at a quantitative level; this requires further investigation.

Curiously however, clashes appeared to increase for mRNA genes, including essential genes; it must however be noted that the expression levels of mRNA genes would only be a fraction of rRNA levels. Therefore, it appears that any suppression in the growth defect may arise from a reversal of increased replication-transcription conflicts at rRNA loci, notwithstanding any effect on essential or non-essential mRNA genes.

## Discussion

Taken together, our results indicate that under *ΔrnhA-ΔdnaA* cSDR, selection favours preferential replication initiation from *oriK45*, located ∼0.4-0.7 Mb clockwise of *oriC*. *oriK45* is a broadly-defined region, and spans an ∼300 kb region across the samples analysed here. The precise location of one or more initiation sites within *oriK45* is unknown, and may be beyond the capabilities of MFA experiments in unsynchronised populations. The top homology to R-loop-forming sequences is at ∼4.51 Mb; *hotH*, previously shown to be capable of autonomous replication (28), is ∼40-50 kb clockwise of the above R-loop-forming sequence; *fimD*, an insertion in which has an effect on *ΔtopA*-mediated cSDR (36), is located between the above two sites and is closer to *hotH*. These three sites, while located in the broad region that defines *oriK45*, do not overlap. There could be multiple discrete initiation sites within *oriK45*, or *oriK45* might encompass a region with diffuse initiation points.

Are there one or more *oriK* sites within *ter*? Though the *ter* peak reported by Maduike et al. and Dimude et al. (23, 27) disappeared in a *tus* mutant (as well as in a mutant carrying a small deletion in *tus* in our study), which accounts for fork trapping, this evidence may not fully eliminate the possibility of a relatively weak *ter oriK*. The absence of a strong *ori-ter* gradient in these slow growing *tus* mutants may always cause such a peak to be missed. Though we observe that the *ter* peak is retained in our stationary phase cells, there is still the possibility that there is still some cSDR activity from a *ter* proximal *oriK* site in these cells. That said however, *oriK45* appears to be favoured by selection, and the fact that this site is located relatively close to the canonical *oriC* may help its cause. One potential future experiment would be an analysis of a strain that combines the inversion-containing suppressors isolated in our study with a *Δtus* mutation. In such a mutant, a strong *oriK* site within *ter* might manifest as an obvious peak, but might come with the cost of dramatically upsetting the highly favourable copy number gradient declining from *oriC* towards *ter.*Replication initiation from *oriK45* would result in head-on collisions with RNA polymerases transcribing four rRNA operons encoded between *oriC* and *oriK45*. Such head-on collisions are detrimental at least in part because of DNA topological issues that cause excessive R-loop formation in such conflict sites (42) The predominant suppressor found here would invert the DNA around *oriC* such that these four rRNA operons would now be on the leading strand of replication from *oriK45*. This would however place one rRNA operon now on the lagging strand. The promoter of this rRNA operon carried a mutation in the discriminator region in all inversion-carrying suppressor strains. Though we couldn’t find any significant difference in the expression levels of plasmid-borne GFP cloned downstream of the wildtype *rrnD* promoter and that with the discriminator mutation (Supplementary figure 4), whether this mutation confers a specific ppGpp-dependent effect on gene expression in a cSDR background, and whether this affects fitness remains to be understood. Recent evidence shows that certain genes – including determinants of virulence and antibiotic resistance – over the course of evolution might have switched in the reverse direction: from leading to lagging (43). Such genes experience higher rates of non-synonymous mutations, experiencing positive selection and thereby promoting evolvability. However, this would not apply to highly expressed genes such as the rRNA genes.

In a previous study, the Sherratt lab placed a second *ori* termed *oriZ* ∼1 Mb clockwise of *oriC*. They reported that replication initiation from *oriZ*, despite *oriZ* being positioned such that it would cause replication-transcription conflicts at rRNA operons, caused little replication or growth defects (44). However, a later attempt by Ivanova and colleagues to create a similar strain revealed a strong growth defect, and also showed that mutations that allow the RNA polymerase to bypass conflicts efficiently, and those that inactivate *ter* can suppress the growth defect (45). MFA analysis of the Sherratt lab strain by Ivanova et al. indicated the presence of a large inversion, affecting several rRNA operons, which had not been detected by the Sherratt study (44). The inversion reported by Ivanova et al. (45) is similar to that observed in our study, except that the right end reported by the earlier study extends beyond that found by us to a position closer to that of *oriZ*. Thus, Ivanova et al. could conclude that replication-transcription conflicts are key determinants of fitness of *E. coli*. These findings are consistent with those of Srivatsan et al. who showed that a large *oriC*-proximal inversion can cause growth defects when *Bacillus subtilis* is grown in rich media (8). Contrary to these findings, Esnault et al. showed that inversions near *oriC* which would place 1-3 rRNA operons on the lagging strand of replication, showed little growth defect (41). That the inversion observed in our study contributes to fitness may be ascertained from the fact that the larger inversion produces higher copy number gradients than the smaller inversion, although both strains carry the *rrnD* promoter mutation. The selective advantage conferred by the inversion also indicates that replication initiates predominantly clockwise of *oriC*, from a position that is also clockwise of the four rRNA operons that are inverted. *oriK45* satisfies these requirements.

Structural variations around *ter* have also been found to exist in *E. coli* with a second *ori*. Dimude et al. placed a second *ori*, termed *oriX,* counterclockwise of *oriC*. They found that this mutant carried a ∼0.8 Mb inversion spanning the *ter* (46). However, this mutant grew slowly. Since the authors did not isolate an *oriX*+ strain without the inversion, they were unable to directly test whether it conferred a selective advantage, even if a small one, to its parent.

Whereas the previous studies by Ivanova et al., and Dimude et al., (45, 46) isolated structural variations while making the parent strain, we were able to isolate our suppressors only after 4-8 days of selection in a laboratory evolution experiment.

Though cSDR may not necessarily be a physiological or natural phenomenon in *E. coli*, with the possible exception of its manifestation as nSDR in stationary phase, it has been argued that this could be a potential primordial mechanism of DNA replication initiation (11). Further, cSDR can provide the bacterium avenues for the development of resistance against new antibiotics targeting initiation of DNA replication (47, 48).

## Materials and methods

### Strains and Media conditions

Wild type (*rnhA*+*dnaA*+) strain mentioned in this study is a derivative of non pathogenic *E.coli* K12 MG1655 strain named GJ13519 in (30). Gene deletions were performed using the one-step inactivation method described by Datsenko and Wanner (49) or by P1 phage mediated transduction (50). All experiments were conducted in Luria Bertani (LB; Hi-Media, India, M575-500) broth to select for suppressors at a faster rate compared to a slow growing minimal media conditions used in previous studies of cSDR. Higher growth rates also produce stronger ori-ter gradients, which enable better peak identification. Where required, the strains were grown in the presence of antibiotics Kanamycin, Ampicillin and Trimethoprim at a final concentrations of 50, 50 and 10 μg/ml respectively.

Growth curves were generated in 250ml flasks or 24-well plates in Luria Bertani (LB; Hi-Media, India, M575-500) broth at 37°C with shaking at 200 rpm. Optical density (OD) measurements were carried out at 600 nm (OD 600) using UV-visible spectrophotometer (SP-8001) or multi well plate reader (Infinite F200pro, Tecan). Growth rates were calculated using Growthcurver (https://CRAN.R-project.org/package=growthcurver) and all plots were generated using customized R scripts.

### Spotting Assay

Spotting assay was performed for all strains at μmax, which corresponds to the maximum growth at the exponential phase of growth. Overnight grown bacterial cultures were diluted in LB media to achieve 0.03 OD and incubated at 37 °C, 200 rpm until μmax. Serial 10-fold dilutions of cultures were spotted (as 3 μl spots) on LB agar plates. The plates were imaged after 30 hours of incubation at 37 °C.

### Whole genome sequencing and DNA copy number analysis

For genomic DNA extraction, the overnight cultures were inoculated in 50 ml of fresh LB media to bring the initial Optical Density (OD) of the culture to 0.03 and the flasks were incubated at 37°C with shaking at 200 rpm. Cells were harvested at μmax and genomic DNA was isolated using GenEluteTM Bacterial Genomic DNA Kit (NA2120-1KT, Sigma-Aldrich) using the manufacturer’s protocol. Library preparation was carried out using Truseq Nano DNA low throughput Library preparation kit (15041757) and Paired end (2X100) sequencing of genomic DNA was performed on the Illumina Hiseq 2500 platform. For stationary phase whole genome sequencing, the cultures were harvested after 16 hours of growth.

The sequencing reads were aligned and mapped to the reference genome (NC_000913.3) using Burrows Wheeler Aligner (BWA) (51) specifying alignment quality and mapping quality thresholds as 20. Read coverage across the genome was calculated for non-overlapping windows of 200nt each using custom PERL scripts and the values were normalized by the mode of the distribution across these bins. The normalized values in logarithmic scale (log2) were plotted against chromosome coordinates to get measures of DNA copy number from *ori* to *ter.* The coordinates were repositioned in such a way that the numbering starts from *oriC* position in either direction. Loess polynomial regression analysis was used for curve fitting.

### Laboratory Evolution of cSDR mutant

Laboratory evolution experiment was carried out for overnight grown cultures of eight independent isolates of *ΔrnhA-ΔdnaA*. Cells were grown in 24-well plates at 37°C, shaking at 200 rpm, until late exponential phase and diluted by a factor of 1:100 into fresh LB broth. Bacterial populations were stored as 50% glycerol stocks at -80 degree Celsius before the next sub-culturing. Contamination check was done for each population using PCR amplification of *rnhA* and *dnaA* genes from isolated genomic DNA samples. Alternative passages were plated on Luria agar plates (10-6 and 10-7 dilution) and counted CFU/ml for each sample during the course of evolution. Number of generations of evolution (N) was calculated using the minimum and maximum OD values per passage. The growth characteristics of evolved populations were monitored in 96-well plates at 37°C, 200 rpm using a Plate reader (Tecan, infinite® F200 PRO). Randomly chosen colonies from different passages were selected for whole genome sequencing.

### Mutation analysis and ori-to-ter ratio calculation

SNPs and indels were identified from the genome sequencing data using the BRESEQ (version 0.33.1) (52) pipeline which uses Bowtie for sequence alignment. A mutational matrix representing presence and absence of mutations were generated from BRESEQ output file using custom R scripts and heat maps were generated using Matrix2png (53). Copy number plots for each sample at the maximum growth rate were used to determine *ori*-to-*ter* ratios. The ratio of maximum loess fit value (excluding *ter*) to the loess fit value of *dif* site (1588800) for each evolved strain was calculated using custom scripts.

### Nanopore sequencing and assembly of genomes

Genomic DNA isolated using GenElute^TM^ Bacterial Genomic DNA Kit (NA2120-1KT, Sigma Aldrich) were subjected to Nanopore sequencing. Sequencing library preparation was carried out with Nanopore Genomic Sequencing Kit SQK-108 and a PCR-free ‘native barcoding’ kit following manufacturer’s protocol. Barcoded samples were pooled and loaded onto a MinION^TM^ Flow Cell MIN106 controlled by MinKNOW version V1.2.8 software (ONT). Base calling was performed using albacore Basecall_Barcoding workflow (version 1.11.5) (ONT). The Fasta files of reads obtained from sequencing were subjected to a Denovo assembler Canu (https://github.com/marbl/canu) using default parameters. Assembled contigs were analysed using sequence aligner Mauve (http://darlinglab.org/mauve/mauve.html) to find chromosomal rearrangements.

### *OriC* inversion prediction in *E.coli* genomes

675 complete *E.coli* genomes downloaded from ncbi ftp site were used for this analysis. For finding *E.coli* strains which possess a chromosomal inversion of *oriC* region, blastn was performed on genomes with *E.coli* K12 MG1655 (NC_000913.3) genome as query and reference for inversion. The inverted regions from blast output of complete genomes were stitched and added together to calculate the total inverted region, thus an inference was made on the status of inversion of region involving *oriC*. *oriC* positions in these genomes were predicted for all *E.coli* strains by performing blastn using *E.coli* K12 MG1655 (NC_000913.3) *oriC* region as query.

### *oriK* peak prediction

*oriK* positions were predicted from the loess fitted copy number plots using custom R scripts. Outliers were removed by visualization from the copy number data before fitting the curve. The loess fit was derived after removing known deletions and reversing the copy number curve around inversions. A position was called as an *oriK* peak if it has a negative slope, measured relative to the peak position, up to 100 kbp in both directions in the loess predicted values. Peak range (mentioned in Supplementary table 5) is defined from the minimum to maximum position predicted for each peak site across strains.

### R-loop predictions using QmRLFs Finder

To predict RNA-DNA hybrids on the chromosome we used QmRLFs model (34, 35) on *Escherichia coli* K12 MG1655 (NC_000913.3) genome with default parameters. From the output file we considered starting position of a predicted R-loop and plotted a line plot for these positions using custom R scripts for both the models (m1 and m2) separately.

### RNA extraction, mRNA enrichment and sequencing

Overnight cultures were inoculated in 100 ml of fresh LB media to bring the initial Optical Density (OD) of the culture to 0.03 and the flasks were incubated at 37°C with shaking at 200 rpm. Samples were collected at the maximum growth rate and two biological replicates were performed for each sample. The samples were immediately processed for total RNA isolation using Trizol method (15596018; Invitrogen). DNase treated RNA was depleted of ribosomal RNA using the Ambion MicrobeExpressTM Kit (AM1905). Libraries were prepared for RNA-sequencing using NEBNextUltra Directional RNA Library Prep Kit for Illumina (New England Biolabs), according to manufacturer’s protocol and single end sequencing for 50 cycles were done using Illumina HiSeq 2500 platform.

### Transcriptome analysis

The sequencing reads were aligned and mapped to the reference genome (NC_000913.3) using Burrows Wheeler Aligner (BWA) (51). The reference genome sequence (.fna) and annotation (.gff) files for the same strain were downloaded from the ncbi ftp website (ftp://ftp.ncbi.nlm.nih.gov ). The raw read quality was checked using the FastQC software (version v0.11.5). SAMTOOLS (version 1.2) and BEDTOOLS (verson 2.25.0) were used to calculate the read count per gene using the annotation file (.bed). The format of the annotation file (.gff) was changed to .bed using an in-house python script. The normalization and differential gene expression analysis for the two conditions were carried out using the edgeR pipeline (54). Log fold change expression values in comparison to *ΔrnhA-ΔdnaA* were plotted using In-house R scripts and the pearson correlation values were predicted for the same. The genes that are differentially expressed by a log(base 2) fold change of 1.5 or above with FDR value of 0.01 were considered as differentially expressed.

### DNA Copy number and transcriptome comparison

The sequencing reads of both DNA and RNA isolated at the exponential phase of growth were analysed as similar to transcriptome analysis described in materials and methods. The normalization and differential gene expression analysis for the two conditions were carried out using the edgeR (54) pipeline. Smoothed LogFC expression values in comparison to *ΔrnhA-ΔdnaA* were plotted against chromosome coordinates using in-house R scripts.

### Probability of head-on collision prediction

The probability of head-on collisions in evolved and parental strains from RNA sequencing data was calculated for the chromosome region 3.3Mb to 4.6Mb, which includes the inversion. The rate of head-on collisions in the presence or absence of the inversion was calculated by assuming the activation of a single predominant origin of replication in evolved and parental clones (either *oriC* or *oriK45*). The fractional score of head-on replication-transcription conflicts was defined as the ratio of the number of reads mapping to genes encoded on the lagging strand to the total number of reads mapping to the region for each strain. The strand information for genes was adapted from NC_000913 (version 3) .ptt or .rnt files.

### Promoter activity assay

The promoter activity of the mutant and wild type *rrnD* promoter region (rrsD-yrdA Intergenic region (IGR)) was monitored by transforming the pUA139 plasmid containing cloned construct of the IGR region in to wild type *E.coli*. M9 medium with 0.2% glucose was used to culture the strains. Overnight culture containing the plasmid strain was inoculated at a ratio of 1:100 in 100 ml media and the samples were isolated at various intervals to measure GFP fluorescence using FACS caliber. Around 25,000 cells were acquired for each sample using a 488-nm excitation laser, and the emission was recorded from FL1 channel that uses a 530/30 band-pass (BP) filter to collect the GFP intensity pUA139::*gfp plasmid* was used to set the background fluorescence, and GFP intensity above this background was marked as positive. Data analysed using FlowJo software.

## Acknowledgments

We thank Prof. J. Gowrishankar for discussions, materials and comments on our manuscript. We thank Dr. Nalini Raghunathan, Sayantan Goswami and all Gowrishankar lab members at CDFD for discussions and materials. We thank Mohak Sharda for providing DnaA sequence homology data and all Aswin lab members for discussions and comments. We thank Dr. Sunil Laxman (Instem, Bangalore), Prof. Dasaradhi Palakodeti (Instem, Bangalore), Prof. P V Shivaprasad (NCBS-TIFR, Bangalore), Dr. Anjana Badrinarayan (NCBS-TIFR, Bangalore) and Prof. Sandeep Krishna (NCBS-TIFR, Bangalore) for discussions and comments. We acknowledge Dr Awadhesh Pandit and Tejali Naik at the Next Generation Genomics Facility, NCBS for providing sequencing services and CIFF facility at NCBS for technical support.

## Data availability

The genome sequence data from this work are available at https://www.ncbi.nlm.nih.gov/bioproject/PRJNA562391.

The RNA sequence data and processed files from this work are available at https://www.ncbi.nlm.nih.gov/geo/query/acc.cgi?acc=GSE135706.

## Supplementary Figure legends

**Figure S1:** Δ*r*nhA*-ΔdnaA* shows reduced growth in LB media. (A) growth curves of *rnhA^+^dnaA^+^*, *ΔrnhA*, and *ΔrnhA-ΔdnaA* in LB at 37°C, 200 rpm. X-axis indicates time and Y-axis indicates log_2_ OD600. (B) and (C) box plots for lag time and growth rate followed by each strain respectively. *ΔrnhA-ΔdnaA* shows reduced growth rate and extended lag phase compared to *rnhA^+^dnaA^+^* (P<<<-1, Wilcoxon test, one-tailed). (D) Spotting assay for *rnhA^+^dnaA^+^, ΔrnhA*, and *ΔrnhA-ΔdnaA* using different dilutions of cultures (left to right: 10^-3^, 10^-4^ , 10^-5^ and 10^-6^) in Luria agar plates incubated at 37°C.

**Figure S2:** Plot represents mutational matrices representing pseudogene mutations and common mutations (which are present in parental strains) for independent colonies sequenced. Red colour indicates the presence of the mutation in the gene shown on Y-axis. X-axis shows the sample IDs of suppressor mutants and parental strains evolved from three independent populations.

**Figure S3:** Plots showing the trend followed by the smoothened log_2_fold change values for all genes in comparison with *ΔrnhA-ΔdnaA* strain for gene expression and DNA copy number. (A) *rnhA^+^dnaA^+^/ΔrnhA-ΔdnaA*, (B) *ΔrnhA/ΔrnhA-ΔdnaA,* (C) *ΔrnhA-ΔdnaAinv^rrnD-rrnE^/ΔrnhA-dnaA*, and (D) *ΔrnhA-ΔdnaAinv^rrnD-rrnC^/ΔrnhA-ΔdnaA* . X-axis represents positions centered around *oriC* and Y axis represents loess fit values of log_2_ fold change. Red lines represent gene expression and black lines represent DNA copy number for the same strain.

**Figure S4:** Plots showing the respective GFP fluorescence intensity measured using FACS for *pUA139* (orange), *wt_rrnD_IGR_pUA139* (blue) and *mut_rrnD_IGR_pUA139* (red) strains at different time intervals, (A) 0 hour (B) 2hours and (C) 5hours.

## Supplementary Table legends

**Table S1:** List of eubacterial strains which did not have a *dnaA* homologue detected in their genomes (out of 5976 organisms).

**Table S2:** *Ori-to-ter* ratios for all strains calculated from MFA plots. To read the sample ids, first number represents the evolution lane(1,5,8), followed by D letter and the number which represents Day of evolution(0, 4, 8, 12, 15), and last number represents the number of colony selected from that population. For example 5D0_3 means 3^rd^ colony selected from the 0^th^ day of evolution of evolution lane number 5.

**Table S3:** List of *E.coli* strains in which presence of a chromosomal inversion around *oriC* is observed.

**Table S4:** Chromosomal position of predicted *oriK* peaks from MFA plots. The value represents loess fit value for the maxima position of the peak identified.

**Table S5:** Functional classification of genes based on COG categories. Enrichment of a functional category is marked in yellow.

**Table S6:** Strains, plasmids and primers used in this study.

## References

1. Mott ML, Berger JM. 2007. DNA replication initiation: mechanisms and regulation in bacteria. Nat Rev Microbiol 5:343.

2. Duggin IG, Bell SD. 2009. Termination Structures in the Escherichia coli Chromosome Replication Fork Trap. J Mol Biol 387:532–539.

3. Rocha EP. 2004. The replication-related organization of bacterial genomes. Microbiology 150:1609–1627.

4. Couturier E, Rocha EP. 2006. Replication-associated gene dosage effects shape the genomes of fast-growing bacteria but only for transcription and translation genes. Mol Microbiol 59:1506– 1518.

5. Khedkar S, Seshasayee ASN. 2016. Comparative Genomics of Interreplichore Translocations in Bacteria: A Measure of Chromosome Topology? G3 Genes Genomes Genet 6:1597–1606.

6. Bryant JA, Sellars LE, Busby SJ, Lee DJ. 2014. Chromosome position effects on gene expression in Escherichia coli K-12. Nucleic Acids Res 42:11383–11392.

7. Wang JD, Berkmen MB, Grossman AD. 2007. Genome-wide coorientation of replication and transcription reduces adverse effects on replication in Bacillus subtilis. Proc Natl Acad Sci 104:5608–5613.

8. Srivatsan A, Tehranchi A, MacAlpine DM, Wang JD. 2010. Co-orientation of replication and transcription preserves genome integrity. PLoS Genet 6:e1000810.

9. Clayton DA. 1982. Replication of animal mitochondrial DNA. Cell 28:693–705.

10. Yasukawa T, Kang D. 2018. An overview of mammalian mitochondrial DNA replication mechanisms. J Biochem (Tokyo) 164:183–193.

11. Kogoma T. 1997. Stable DNA replication: interplay between DNA replication, homologous recombination, and transcription. Microbiol Mol Biol Rev 61:212–238.

12. Kogoma T, Torrey TA, Connaughton MJ. 1979. Induction of UV-resistant DNA replication in Escherichia coli: Induced stable DNA replication as an SOS function. Mol Gen Genet MGG 176:1–9.

13. Ogawa T, Pickett GG, Kogoma T, Kornberg A. 1984. RNase H confers specificity in the dnaA-dependent initiation of replication at the unique origin of the Escherichia coli chromosome in vivo and in vitro. Proc Natl Acad Sci 81:1040–1044.

14. Martel M, Balleydier A, Sauriol A, Drolet M. 2015. Constitutive stable DNA replication in Escherichia coli cells lacking type 1A topoisomerase activity. DNA Repair 35:37–47.

15. Gowrishankar J, Harinarayanan R. 2004. Why is transcription coupled to translation in bacteria? Mol Microbiol 54:598–603.

16. Gowrishankar J, Leela JK, Anupama K. 2013. R-loops in bacterial transcription: their causes and consequences. Transcription 4:153–157.

17. Harinarayanan R, Gowrishankar J. 2003. Host factor titration by chromosomal R-loops as a mechanism for runaway plasmid replication in transcription termination-defective mutants of Escherichia coli. J Mol Biol 332:31–46.

18. Raghunathan N, Kapshikar RM, Leela JK, Mallikarjun J, Bouloc P, Gowrishankar J. 2018. Genome-wide relationship between R-loop formation and antisense transcription in Escherichia coli. Nucleic Acids Res 46:3400–3411.

19. Hong X, Cadwell GW, Kogoma T. 1995. Escherichia coli RecG and RecA proteins in R-loop formation. EMBO J 14:2385–2392.

20. Lloyd RG, Rudolph CJ. 2016. 25 years on and no end in sight: a perspective on the role of RecG protein. Curr Genet 62:827–840.

21. Rudolph CJ, Upton AL, Stockum A, Nieduszynski CA, Lloyd RG. 2013. Avoiding chromosome pathology when replication forks collide. Nature 500:608–611.

22. Midgley-Smith SL, Dimude JU, Taylor T, Forrester NM, Upton AL, Lloyd RG, Rudolph CJ. 2018. Chromosomal over-replication in Escherichia coli recG cells is triggered by replication fork fusion and amplified if replichore symmetry is disturbed. Nucleic Acids Res 46:7701– 7715.

23. Dimude JU, Stockum A, Midgley-Smith SL, Upton AL, Foster HA, Khan A, Saunders NJ, Retkute R, Rudolph CJ. 2015. The consequences of replicating in the wrong orientation: bacterial chromosome duplication without an active replication origin. MBio 6:e01294–15.

24. Raghunathan N, Goswami S, Leela JK, Pandiyan A, Gowrishankar J. 2019. A new role for Escherichia coli Dam DNA methylase in prevention of aberrant chromosomal replication. Nucleic Acids Res 47:5698–5711.

25. Hong X, Cadwell GW, Kogoma T. 1996. Activation of stable DNA replication in rapidly growing Escherichia coli at the time of entry to stationary phase. Mol Microbiol 21:953–961.

26. de Massy B, Fayet O, Kogoma T. 1984. Multiple origin usage for DNA replication in sdrA (rnh) mutants of Escherichia coli K-12: initiation in the absence of oriC. J Mol Biol 178:227– 236.

27. Maduike NZ, Tehranchi AK, Wang JD, Kreuzer KN. 2014. Replication of the E scherichia coli chromosome in RN ase HI-deficient cells: multiple initiation regions and fork dynamics. Mol Microbiol 91:39–56.

28. Nishitani H, Hidaka M, Horiuchi T. 1993. Specific chromosomal sites enhancing homologous recombination in Escherichia coli mutants defective in RNase H. Mol Gen Genet MGG 240:307–314.

29. Gowrishankar J. 2015. End of the beginning: elongation and termination features of alternative modes of chromosomal replication initiation in bacteria. PLoS Genet 11:e1004909.

30. Leela JK, Syeda AH, Anupama K, Gowrishankar J. 2013. Rho-dependent transcription termination is essential to prevent excessive genome-wide R-loops in Escherichia coli. Proc Natl Acad Sci 110:258–263.

31. Brochu J, Vlachos-Breton É, Sutherland S, Martel M, Drolet M. 2018. Topoisomerases I and III inhibit R-loop formation to prevent unregulated replication in the chromosomal Ter region of Escherichia coli. PLoS Genet 14:e1007668.

32. Skovgaard O, Bak M, Løbner-Olesen A, Tommerup N. 2011. Genome-wide detection of chromosomal rearrangements, indels, and mutations in circular chromosomes by short read sequencing. Genome Res 21:1388–1393.

33. Hill CW, Harnish BW. 1981. Inversions between ribosomal RNA genes of Escherichia coli. Proc Natl Acad Sci U S A 78:7069–7072.

34. Jenjaroenpun P, Wongsurawat T, Yenamandra SP, Kuznetsov VA. 2015. QmRLFS-finder: a model, web server and stand-alone tool for prediction and analysis of R-loop forming sequences. Nucleic Acids Res 43:10081.

35. Kuznetsov VA, Bondarenko V, Wongsurawat T, Yenamandra SP, Jenjaroenpun P. 2018. Toward predictive R-loop computational biology: genome-scale prediction of R-loops reveals their association with complex promoter structures, G-quadruplexes and transcriptionally active enhancers. Nucleic Acids Res 46:7566–7585.

36. Usongo V, Martel M, Balleydier A, Drolet M. 2016. Mutations reducing replication from R-loops suppress the defects of growth, chromosome segregation and DNA supercoiling in cells lacking topoisomerase I and RNase HI activity. DNA Repair 40:1–17.

37. Dai Y, Outten FW. 2012. The E. coli SufS-SufE sulfur transfer system is more resistant to oxidative stress than IscS-IscU. FEBS Lett 586:4016–4022.

38. Viguera E, Petranovic M, Zahradka D, Germain K, Ehrlich DS, Michel B. 2003. Lethality of bypass polymerases in Escherichia coli cells with a defective clamp loader complex of DNA polymerase III. Mol Microbiol 50:193–204.

39. Valens M, Penaud S, Rossignol M, Cornet F, Boccard F. 2004. Macrodomain organization of the Escherichia coli chromosome. EMBO J 23:4330–4341.

40. Lal A, Dhar A, Trostel A, Kouzine F, Seshasayee ASN, Adhya S. 2016. Genome scale patterns of supercoiling in a bacterial chromosome. Nat Commun 7:11055.

41. Esnault E, Valens M, Espéli O, Boccard F. 2007. Chromosome structuring limits genome plasticity in Escherichia coli. PLoS Genet 3:e226.

42. Lang KS, Merrikh H. 2019. Topological stress is responsible for the detrimental outcomes of head-on replication-transcription conflicts. bioRxiv 691188.

43. Merrikh CN, Merrikh H. 2018. Gene inversion potentiates bacterial evolvability and virulence. Nat Commun 9:4662.

44. Wang X, Lesterlin C, Reyes-Lamothe R, Ball G, Sherratt DJ. 2011. Replication and segregation of an Escherichia coli chromosome with two replication origins. Proc Natl Acad Sci U S A 108:E243–250.

45. Ivanova D, Taylor T, Smith SL, Dimude JU, Upton AL, Mehrjouy MM, Skovgaard O, Sherratt DJ, Retkute R, Rudolph CJ. 2015. Shaping the landscape of the Escherichia coli chromosome: replication-transcription encounters in cells with an ectopic replication origin. Nucleic Acids Res 43:7865–7877.

46. Dimude JU, Stein M, Andrzejewska EE, Khalifa MS, Gajdosova A, Retkute R, Skovgaard O, Rudolph CJ. 2018. Origins Left, Right, and Centre: Increasing the Number of Initiation Sites in the Escherichia coli Chromosome. Genes 9.

47. Grimwade JE, Leonard AC. 2017. Targeting the Bacterial Orisome in the Search for New Antibiotics. Front Microbiol 8.

48. van Eijk E, Wittekoek B, Kuijper EJ, Smits WK. 2017. DNA replication proteins as potential targets for antimicrobials in drug-resistant bacterial pathogens. J Antimicrob Chemother 72:1275–1284.

49. Datsenko KA, Wanner BL. 2000. One-step inactivation of chromosomal genes in Escherichia coli K-12 using PCR products. Proc Natl Acad Sci U S A 97:6640–6645.

50. Thomason LC, Costantino N, Court DL. 2007. E. coli genome manipulation by P1 transduction. Curr Protoc Mol Biol Chapter 1:Unit 1.17.

51. Li H, Durbin R. 2009. Fast and accurate short read alignment with Burrows-Wheeler transform. Bioinforma Oxf Engl 25:1754–1760.

52. Deatherage DE, Barrick JE. 2014. Identification of mutations in laboratory-evolved microbes from next-generation sequencing data using breseq. Methods Mol Biol Clifton NJ 1151:165– 188.

53. Pavlidis P, Noble WS. 2003. Matrix2png: a utility for visualizing matrix data. Bioinformatics 19:295–296.

54. McCarthy DJ, Chen Y, Smyth GK. 2012. Differential expression analysis of multifactor RNA-Seq experiments with respect to biological variation. Nucleic Acids Res 40:4288–4297.

