## Supplemental Table 1 for "Laboratory Evolution Experiments Help Identify a Predominant Region of Constitutive Stable DNA Replication Initiation"

| | |
|---|---|
| GCF_000008885 | Wigglesworthia_glossinidia_endosymbiont_of_Glossina_brevipalpis_plasmid_pWb1_DNA |
| GCF_000011745 | Candidatus_Blochmannia_pennsylvanicus_str._BPEN |
| GCF_000013185 | Baumannia_cicadellinicola_str._Hc_(Homalodisca_coagulata) |
| GCF_000022605 | Blattabacterium_sp._(Blattella_germanica)_str._Bge_plasmid_pBge |
| GCF_000025125 | Candidatus_Atelocyanobacterium_thalassa_isolate_ALOHA |
| GCF_000043285 | Blochmannia_floridanus_complete_genome |
| GCF_000093065 | Candidatus_Riesia_pediculicola_USDA_plasmid_pPAN |
| GCF_000146025 | Uncultured_Termite_group_1_bacterium_phylotype_Rs-D17_plasmid pTGRD3 DNA |
| GCF_000165505 | Ilyobacter_polytropus_DSM_2926_plasmid_pILYOP02 |
| GCF_000177535 | Corynebacterium_resistens_DSM_45100 |
| GCF_000179035 | Mycoplasma_suis_str._Illinois |
| GCF_000185985 | Candidatus_Blochmannia_vafer_str._BVAF |
| GCF_000196515 | 'Nostoc_azollae'_0708_plasmid_pAzo02 |
| GCF_000203215 | Mycoplasma_suis_KI3806_complete_genome |
| GCF_000219175 | Candidatus_Moranella_endobia_PCIT |
| GCF_000223375 | Ketogulonigenium_vulgarum_WSH-001_plasmid_2 |
| GCF_000236405 | Blattabacterium_sp._(Cryptocercus_punctulatus)_str._Cpu_plasmid_pCpu |
| GCF_000247565 | Wigglesworthia_glossinidia_endosymbiont_of_Glossina_morsitans_morsitans |
| GCF_000255275 | Corynebacterium_diphtheriae_PW8 |
| GCF_000262655 | Helicobacter_pylori_XZ274_plasmid_pXZ274 |
| GCF_000287295 | Candidatus_Carsonella_ruddii_HT_isolate_Thao2000 |
| GCF_000292685 | Candidatus_Portiera_aleyrodidarum_BT-B |
| GCF_000298385 | Candidatus_Portiera_aleyrodidarum_BT-QVLC |
| GCF_000300035 | Candidatus_Portiera_aleyrodidarum_BT-QVLC |
| GCF_000300075 | Candidatus_Portiera_aleyrodidarum_BT-B |

| | |
|---|---|
| GCF_000304735 | Borrelia_afzelii_HLJ01 |
| GCF_000317675 | Cyanobacterium_aponinum_PCC_10605_plasmid_pCYAN10605.01 |
| GCF_000319385 | Candidatus_Endolissoclinum_faulkneri_L2 |
| GCF_000331065 | Candidatus_Blochmannia_chromaiodes_str._640 |
| GCF_000364725 | Candidatus_Moranella_endobia_PCVAL |
| GCF_000441555 | Candidatus_Profftella_armatura_plasmid |
| GCF_000471965 | Blattabacterium_sp._(Nauphoeta_cinerea)_plasmid |
| GCF_000477415 | Mycoplasma_parvum_str._Indiana |
| GCF_000505725 | Francisella_noatunensis_subsp._orientalis_LADL--07-285A |
| GCF_000508245 | Mycoplasma_ovis_str._Michigan |
| GCF_000604125 | Treponema_pallidum_subsp._pallidum_str._Sea_81-4 |
| GCF_000709555 | Endosymbiont_of_Llaveia_axin_axin |
| GCF_000767685 | Corynebacterium_ulcerans_FRC11 |
| GCF_000769635 | Corynebacterium_ulcerans_strain_05146 |
| GCF_000770175 | Mycobacterium_abscessus_strain_DJO-44274 |
| GCF_000815025 | Coxiella_endosymbiont_of_Amblyomma_americanum |
| GCF_000827855 | Candidatus_Portiera_aleyrodidarum_MED_(Bemisia_tabaci)_strain_BT-Q |
| GCF_000828815 | Candidatus_Tachikawaea_gelatinosa_DNA |
| GCF_000828835 | Thioploca_ingrica_DNA |
| GCF_000829235 | Cyanobacterium_endosymbiont_of_Epithemia_turgida_isolate_EtSB_Lake_Yunoko_DNA |
| GCF_000953435 | Candidatus_Evansia_muelleri_genome_assembly_CEM1.1 |
| GCF_000973545 | Blochmannia_endosymbiont_of_Camponotus_(Colobopsis)_obliquus_strain_757 |
| GCF_001021025 | Corynebacterium_epidermidicanis_strain_DSM_45586 |
| GCF_001278785 | Candidatus_Profftella_armatura_strain_YCPA_plasmid |
| GCF_001318295 | Candidatus_Xiphinematobacter_sp._Idaho_Grape |
| GCF_001548095 | Geminocystis_sp._NIES-3708_plasmid_pGM05_DNA |
| GCF_001548115 | Geminocystis_sp._NIES-3709_plasmid_pGM3709_11_DNA |
| GCF_001587015 | Campylobacter_jejuni_strain_OD267_plasmid_pCJDM67_S |

| | |
|---|---|
| GCF_001682195 | Flammeovirga_sp._MY04_plasmid |
| GCF_900048035 | Enterobacteriaceae_bacterium_symbiont_of_Ferrisia_virgata_isolate_GEFVIR_genome_assembly |
| GCF_900048045 | Enterobacteriaceae_bacterium_symbiont_of_Paracoccus_marginatus_isolate_MEPMAR_genome_assembly |
