## Supplemental Table 2 for "Laboratory Evolution Experiments Help Identify a Predominant Region of Constitutive Stable DNA Replication Initiation"

| Strain ID | ***Ori*-to-*ter* ratio** |
|---|---|
| ***Parental strains*** | |
| 1D0_1 | 1.13 |
| 1D0_2 | 1 |
| 1D0_3 | 1.09 |
| 1D0_4 | 1.09 |
| 5D0_1 | 1.13 |
| 5D0_2 | 1.15 |
| 5D0_3 | 1.07 |
| 5D0_4 | 1.15 |
| 8D0_1 | 1.03 |
| 8D0_2 | 0.97 |
| 8D0_3 | 0.87 |
| 8D0_4 | 1 |
| *ΔrnhA-ΔdnaA* | 1.02 |
| *ΔrnhA* | 1.79 |
| *ΔrnhA-ΔdnaA/pHYD2388* | 1.22 |
| GJ13519 | 2.48 |
| K12 MG1655 | 2.35 |
| ***Suppressor mutants*** | |
| 1D4_1 | 1.02 |
| 1D4_2 | 1.73 |
| 1D4_3 | 1.69 |
| 1D4_4 | 1.03 |
| 1D8_1 | 1.51 |
| 1D8_2 | 1.45 |
| 1D8_3 | 0.94 |

| | |
|---|---|
| 1D8_4 | 1.93 |
| 1D12_2 | 1.28 |
| 1D12_3 | 1.27 |
| 1D12_4 | 1.35 |
| 1D15_1 | 1.42 |
| 1D15_2 | 1.49 |
| 1D15_3 | 1.25 |
| 1D15_4 | 1.47 |
| 5D4_1 | 1.41 |
| 5D4_2 | 1.33 |
| 5D4_3 | 1.38 |
| 5D4_4 | 1.20 |
| 5D8_1 | 1.34 |
| 5D8_2 | 1.38 |
| 5D8_3 | 1.47 |
| 5D8_4 | 1.31 |
| 5D12_1 | 1.68 |
| 5D12_2 | 1.82 |
| 5D12_3 | 2.68 |
| 5D12_4 | 1.67 |
| 5D15_1 | 1.82 |
| 5D15_2 | 1.88 |
| 5D15_3 | 1.89 |
| 5D15_4 | 1.96 |
| 8D4_1 | 1.14 |
| 8D4_2 | 1.02 |
| 8D4_3 | 1.03 |

| | |
|---|---|
| 8D4_4 | 1.07 |
| 8D8_1 | 1.44 |
| 8D8_2 | 1.87 |
| 8D8_3 | 1.76 |
| 8D8_4 | 1.36 |
| 8D15_1 | 1.70 |
| 8D15_2 | 1.70 |
| 8D15_4 | 1.72 |
