## Supplemental Table 3 for "Laboratory Evolution Experiments Help Identify a Predominant Region of Constitutive Stable DNA Replication Initiation"

| | |
|---|---|
| GCF_001021635.2_ASM102163v2 | NZ_CP006834.2 *Escherichia coli* APEC O2-211 chromosome |
| GCF_002220215.1_ASM222021v1 | NZ_CP022393.1 *Escherichia coli* strain E62 chromosome |
| GCF_003052645.1_ASM305264v1 | NZ_CP028702.1 *Escherichia coli* strain J53 chromosome |
| GCF_000499485.1_MYMC4100 | NZ_HG738867.1 *Escherichia coli* str. K-12 substr. MC4100 |
| GCF_003966445.1_ASM396644v1 | NZ_AP018802.1 *Escherichia coli* E2863 DNA |
| GCF_000091005.1_ASM9100v1 | NC_013361.1 *Escherichia coli* O26:H11 str. 11368 DNA |
| GCF_003018715.1_ASM301871v1 | NZ_CP027331.1 *Escherichia coli* strain 2013C-3277 chromosome |
| GCF_002211725.1_ASM221172v1 | NZ_CP022154.1 *Escherichia coli* strain ABWA45 chromosome |
| GCF_003112145.1_ASM311214v1 | NZ_CP028110.1 *Escherichia coli* O121 str. RM8352 chromosome |
| GCF_002879975.1_ASM287997v1 | NZ_CP025747.1 *Escherichia coli* strain ML35 chromosome |
| GCF_001420935.1_ASM142093v1 | NZ_CP013029.1 *Escherichia coli* strain 2012C-4227 |
| GCF_003052665.1_ASM305266v1 | NZ_CP028703.1 *Escherichia coli* strain ME8067 chromosome |
| GCF_002716885.1_ASM271688v1 | NZ_CP015244.1 *Escherichia coli* O91 str. RM7190 chromosome |
| GCF_003018795.1_ASM301879v1 | NZ_CP027352.1 *Escherichia coli* strain 2012C-4606 chromosome |
| GCF_003017805.1_ASM301780v1 | NZ_CP027325.1 *Escherichia coli* strain 2013C-4830 chromosome |
| GCF_002741215.1_ASM274121v1 | NZ_CP024239.1 *Escherichia coli* O15:H11 strain 90-9272 chromosome |
| GCF_001901105.1_ASM190110v1 | NZ_CP010196.1 *Escherichia coli* strain M9 |
| GCF_000010745.1_ASM1074v1 | NC_013353.1 *Escherichia coli* O103:H2 str. 12009 DNA |
| GCF_001901425.1_ASM190142v1 | NZ_CP010240.1 *Escherichia coli* strain C7 |
| GCF_002012025.1_ASM201202v1 | NZ_CP018970.1 *Escherichia coli* strain Ecol_542 chromosome |
| GCF_000010245.2_ASM1024v1 | NC_007779.1 *Escherichia coli* str. K-12 substr. W3110 DNA |
| GCF_003112185.1_ASM311218v1 | NZ_CP028116.1 *Escherichia coli* O26 str. RM8426 chromosome |
| GCF_003966465.1_ASM396646v1 | NZ_AP018808.1 *Escherichia coli* E2865 DNA |

| | |
|---|---|
| GCF_003018495.1_ASM301849v1 | NZ_CP027552.1 *Escherichia coli* strain 2015C-4498 chromosome |
| GCF_000258025.1_ASM25802v1 | NC_017660.1 *Escherichia coli* KO11FL |
| GCF_000148605.1_ASM14860v1 | NC_017632.1 *Escherichia coli* UM146 |
| GCF_001901215.1_ASM190121v1 | NZ_CP010221.1 *Escherichia coli* strain M19 |
| GCF_003018035.1_ASM301803v1 | NZ_CP027390.1 *Escherichia coli* strain 2015C-4944 chromosome |
| GCF_002057355.1_ASM205735v1 | NZ_CP020107.1 *Escherichia coli* strain 13E0767 chromosome |
| GCF_002796445.1_ASM279644v1 | NZ_CP024889.1 *Escherichia coli* strain AR_0019 chromosome |
| GCF_001900535.1_ASM190053v1 | NZ_CP010122.1 *Escherichia coli* strain C5 |
| GCF_002055605.1_ASM205560v1 | NZ_CP020092.1 *Escherichia coli* strain 13E0725 chromosome |
| GCF_003018155.1_ASM301815v1 | NZ_CP027548.1 *Escherichia coli* strain 2014C-3061 chromosome |
| GCF_003019215.1_ASM301921v1 | NZ_CP027766.1 *Escherichia coli* strain 2013C-3342 chromosome |
| GCF_003018895.1_ASM301889v1 | NZ_CP027387.1 *Escherichia coli* strain 2014C-3057 chromosome |
| GCF_003018575.1_ASM301857v1 | NZ_CP027582.1 *Escherichia coli* strain 2013C-4538 chromosome |
| GCF_002057245.1_ASM205724v1 | NZ_CP020106.1 *Escherichia coli* strain 13E0780 chromosome |
