## Supplemental Table 4 for "Laboratory Evolution Experiments Help Identify a Predominant Region of Constitutive Stable DNA Replication Initiation"

| Sample | 3.84-3.89 (*oriC*) | 0.44-0.55 | 1.42-1.52 | 1.88-2.23 | 2.53-2.6 | 2.95-3.36 | 3.63-3.8 |
|---|---|---|---|---|---|---|---|
| **Parental strains** | | | | | | | |
| WT K12 | 2.36 | - | - | - | - | - | - |
| GJ13519 | 2.5 | - | - | - | - | - | - |
| *ΔrnhA* | 1.9 | - | 1.18 | - | - | - | - |
| *ΔdnaA/dnaA* | 2.33 | - | - | - | - | - | - |
| *ΔrnhA-ΔdnaA/dnaA* | - | 1.33 | 1.22 | 1.17 | 1.1 | - | 1.21 |
| *ΔrnhA-ΔdnaA* | - | 1.63 | 1.85 | 1.5 | - | 1.19 | 1.-5 |
| 1D0_1 | - | 1.35 | 1.33 | 1.27 | 1.15 | 1.17 | - |
| 1D0_2 | - | 1.53 | 1.74 | 1.5 | - | 1.21 | 1.-7 |
| 1D0_3 | - | 1.56 | 1.64 | 1.44 | 1.22 | 1.18 | 1.-7 |
| 1D0_4 | - | 1.34 | 1.34 | 1.41 | 1.32 | 1.36 | 1.-8 |
| 5D0_1 | - | 1.47 | 1.49 | 1.37 | 1.2 | 1.18 | 1.-4 |
| 5D0_2 | - | 1.56 | 1.56 | 1.4 | 1.24 | 1.2 | - |
| 5D0_3 | - | 1.63 | 1.73 | - | - | 1.25 | 1.-9 |
| 5D0_4 | - | 1.56 | 1.55 | 1.43 | 1.25 | 1.21 | 1.-7 |
| 8D0_1 | - | 1.33 | 1.47 | 1.26 | 1.11 | 1.-8 | 1.-2 |
| 8D0_2 | - | 1.5 | 1.76 | 1.42 | 1.26 | 1.18 | 1.-4 |
| 8D0_3 | - | 1.31 | 1.68 | 1.31 | 1.-5 | 1.-6 | 1.-2 |
| 8D0_4 | - | 1.48 | 1.7 | 1.42 | - | 1.19 | 1.-5 |
| **Suppressor strains** | | | | | | | |
| 1D4_1 | - | 1.22 | 1.41 | 1.35 | 1.31 | 1.35 | 1.25 |
| 1D4_4 | - | - | 1.6 | 1.51 | - | - | 1.-4 |
| 1D8_3 | - | - | 1.83 | 1.-9 | - | 1.24 | 1.11 |
| 5D4_1 | - | - | 1.3 | 1.18 | - | - | - |
| 5D4_2 | - | - | 1.28 | 1.15 | - | - | 1.34 |
| 5D4_3 | - | - | 1.29 | 1.16 | - | - | 1.38 |
| 5D4_4 | - | - | 1.29 | 1.12 | - | - | 1.24 |
| 5D8_1 | - | - | 1.29 | 1.16 | - | - | - |
| 5D8_2 | - | - | 1.28 | 1.15 | - | - | 1.38 |
| 5D8_3 | - | - | 1.32 | 1.19 | - | - | - |

| | | | | | | | |
|---|---|---|---|---|---|---|---|
| 5D8_4 | - | - | 1.27 | 1.15 | - | - | 1.32 |
| 5D12_1 | - | - | 1.51 | 1.-9 | - | - | - |
| 5D12_2 | - | 1.8 | 1.39 | 1.-9 | 1.11 | 1.25 | 1.29 |
| 5D12_3 | - | - | 1.28 | - | - | - | - |
| 5D12_4 | - | - | 1.61 | 1.-7 | - | 1.3 | 1.36 |
| 5D15_1 | - | - | 1.41 | 1.17 | - | - | - |
| 5D15_2 | - | - | 1.44 | 1.19 | - | - | 1.75 |
| 5D15_3 | - | - | 1.42 | 1.2 | - | - | 1.74 |
| 5D15_4 | - | - | 1.43 | 1.25 | - | - | - |
| 8D4_1 | - | - | 1.1 | 1.14 | - | 1.18 | - |
| 8D4_2 | - | 1.2 | 1.38 | 1.36 | 1.36 | - | 1.-6 |
| 8D4_3 | - | - | 1.49 | 1.46 | - | 1.44 | - |
| 8D4_4 | - | - | 1.34 | 1.35 | 1.41 | - | - |
| 1D4_2 | - | - | 1.41 | 1.26 | - | 1.58 | - |
| 1D4_3 | - | - | 1.4 | 1.25 | - | - | - |
| 1D8_1 | - | - | 1.63 | 1.18 | - | - | - |
| 1D8_4 | - | - | - | 1.3 | - | - | - |
| 1D8_2 | - | - | 1.32 | 1.18 | - | - | - |
| 1D12_2 | - | - | 1.23 | 1.16 | - | 1.34 | 1.28 |
| 1D12_3 | - | - | 1.35 | 1.13 | - | - | - |
| 1D12_4 | - | - | 1.39 | 1.18 | - | - | - |
| 1D15_1 | - | - | 1.24 | 1.19 | - | - | 1.38 |
| 1D15_2 | - | - | 1.31 | 1.21 | - | - | - |
| 1D15_3 | - | - | 1.49 | 1.13 | - | - | - |
| 1D15_4 | - | 1.57 | 1.29 | 1.22 | - | - | - |
| 8D8_1 | - | - | 1.54 | 1.1 | - | - | - |
| 8D8_2 | - | - | 1.35 | 1.21 | - | - | - |
| 8D8_3 | - | - | 1.37 | 1.18 | - | - | - |
| 8D8_4 | - | - | 1.55 | 1.-6 | - | - | - |
| 8D15_1 | - | - | 1.37 | 1.14 | - | - | - |
| 8D15_2 | - | - | 1.34 | 1.15 | - | - | - |
| 8D15_4 | - | - | 1.33 | - | - | - | - |
