## Supplemental Table 5 for "Laboratory Evolution Experiments Help Identify a Predominant Region of Constitutive Stable DNA Replication Initiation"

| ΔrnhA Downregulated genes | | | | | |
|---|---|---|---|---|---|
| COG categories | COG & DE | COG but !DE | !COG but DE | !COG&!DE | P-value |
| **CELLULAR PROCESSES AND SIGNALING** | | | | | |
| [D] Cell cycle control, cell division | 1 | 32 | 338 | 3557 | 0.35 |
| [M] Cell wall/ membrane/ envelope biogenesis | 12 | 216 | 327 | 3373 | 0.06 |
| [N] Cell motility | 27 | 84 | 312 | 3505 | 3.94x10-7* |
| [O] Post-translational modification | 17 | 124 | 322 | 3465 | 0.1664 |
| [T] Signal transduction mechanisms | 14 | 164 | 325 | 3425 | 0.89 |
| [U] Intracellular trafficking and secretion | 12 | 115 | 327 | 3474 | 0.74 |
| [V] Defense mechanisms | 3 | 46 | 336 | 3543 | 0.79 |
| **INFORMATION STORAGE AND PROCESSING** | | | | | |
| [J] Translation | 8 | 176 | 331 | 3413 | 0.03 |
| [K] Transcription | 17 | 286 | 322 | 3303 | 0.05 |
| [L] Replication, recombination and repair | 6 | 199 | 333 | 3390 | 0.001 |
| **METABOLISM** | | | | | |
| [C] Energy production and conversion | 42 | 245 | 297 | 3344 | 0.0004* |
| [E] Amino acid transport and metabolism | 34 | 328 | 305 | 3261 | 0.55 |
| [F] Nucleotide transport and metabolism | 8 | 90 | 331 | 3499 | 1 |
| [G] Carbohydrate transport and metabolism | 61 | 308 | 278 | 3281 | 2.48x10-7* |
| [H] Coenzyme transport and metabolism | 4 | 152 | 335 | 3437 | 0.003 |
| [I] Lipid transport and metabolism | 5 | 95 | 334 | 3494 | 0.27 |
| [P] Inorganic ion transport and metabolism | 12 | 203 | 327 | 3386 | 0.1 |
| [Q] Secondary metabolites biosynthesis | 4 | 60 | 335 | 3529 | 0.65 |
| **POORLY CHARACTERIZED** | | | | | |

| | | | | | |
|---|---|---|---|---|---|
| [R] General function prediction only | 29 | 372 | 310 | 3217 | 0.34 |
| [S] Function unknown | 23 | 294 | 316 | 3295 | 0.4 |

| **ΔrnhA Upregulated genes** | | | | | |
|---|---|---|---|---|---|
| COG categories | COG & DE | COG but !DE | !COG but DE | !COG&! DE | P-value |
| **CELLULAR PROCESSES AND SIGNALING** | | | | | |
| [D] Cell cycle control, cell division | 3 | 30 | 345 | 3550 | 1 |
| [M] Cell wall/ membrane/ envelope biogenesis | 10 | 218 | 338 | 3362 | 0.011 |
| [N] Cell motility | 13 | 98 | 335 | 3482 | 0.3 |
| [O] Post-translational modification | 9 | 132 | 339 | 3448 | 0.36 |
| [T] Signal transduction mechanisms | 12 | 166 | 336 | 3414 | 0.34 |
| [U] Intracellular trafficking and secretion | 15 | 112 | 333 | 3468 | 0.26 |
| [V] Defense mechanisms | 1 | 48 | 347 | 3532 | 0.12 |
| **INFORMATION STORAGE AND PROCESSING** | | | | | |
| [J] Translation | 35 | 149 | 313 | 3431 | 1.244X10 -5 |
| [K] Transcription | 32 | 271 | 316 | 3309 | 0.29 |
| [L] Replication, recombination and repair | 20 | 185 | 328 | 3395 | 0.61 |
| **METABOLISM** | | | | | |
| [C] Energy production and conversion | 41 | 246 | 307 | 3334 | 0.0016 |
| [E] Amino acid transport and metabolism | 32 | 330 | 316 | 3250 | 1 |
| [F] Nucleotide transport and metabolism | 9 | 89 | 339 | 3491 | 0.85 |
| [G] Carbohydrate transport and metabolism | 29 | 340 | 319 | 3240 | 0.56 |
| [H] Coenzyme transport and metabolism | 7 | 149 | 341 | 3431 | 0.059 |
| [I] Lipid transport and metabolism | 6 | 94 | 342 | 3486 | 0.37 |
| [P] Inorganic ion transport and metabolism | 27 | 188 | 321 | 3392 | 0.06 |

| | | | | | |
|---|---|---|---|---|---|
| [Q] Secondary metabolites biosynthesis | 6 | 58 | 342 | 3522 | 0.82 |
| **POORLY CHARACTERIZED** | | | | | |
| [R] General function prediction only | 28 | 373 | 320 | 3207 | 0.19 |
| [S] Function unknown | 13 | 304 | 335 | 3276 | 0,00092 |

| **ΔrnhA-ΔdnaA Down-regulated genes** | | | | | |
|---|---|---|---|---|---|
| COG categories | COG & DE | COG but !DE | !COG but DE | !COG&! DE | P-value |
| **CELLULAR PROCESSES AND SIGNALING** | | | | | |
| [D] Cell cycle control, cell division | 2 | 31 | 514 | 3381 | 0.3 |
| [M] Cell wall/ membrane/ envelope biogenesis | 17 | 211 | 499 | 3201 | 0.008 |
| [N] Cell motility | 29 | 82 | 487 | 3330 | 0.00016 |
| [O] Post-translational modification | 24 | 117 | 492 | 3295 | 0.16 |
| [T] Signal transduction mechanisms | 22 | 156 | 494 | 3256 | 0.8 |
| [U] Intracellular trafficking and secretion | 15 | 112 | 501 | 3300 | 0.78 |
| [V] Defense mechanisms | 8 | 41 | 508 | 3371 | 0.52 |
| **INFORMATION STORAGE AND PROCESSING** | | | | | |
| [J] Translation | 10 | 174 | 506 | 3238 | 0.0007 |
| [K] Transcription | 34 | 269 | 482 | 3143 | 0.33 |
| [L] Replication, recombination and repair | 14 | 191 | 502 | 3221 | 0.004 |
| **METABOLISM** | | | | | |
| [C] Energy production and conversion | 62 | 225 | 454 | 3187 | 3.71x10-5 |
| [E] Amino acid transport and metabolism | 46 | 316 | 470 | 3096 | 0.87 |
| [F] Nucleotide transport and metabolism | 11 | 87 | 505 | 3325 | 0.65 |
| [G] Carbohydrate transport and metabolism | 82 | 287 | 434 | 3125 | 3.92x10-7 |
| [H] Coenzyme transport and metabolism | 10 | 146 | 506 | 3266 | 0.01 |
| [I] Lipid transport and | 8 | 92 | 508 | 3320 | 0.135 |

| | | | | | |
|---|---|---|---|---|---|
| metabolism | | | | | |
| [P] Inorganic ion transport and metabolism | 22 | 193 | 494 | 3219 | 0.2 |
| [Q] Secondary metabolites biosynthesis | 6 | 58 | 510 | 3354 | 0.45 |
| **POORLY CHARACTERIZED** | | | | | |
| [R] General function prediction only | 60 | 341 | 456 | 3071 | 0.27 |
| [S] Function unknown | 34 | 283 | 482 | 3129 | 0.19 |

| **ΔrnhA-ΔdnaA Up-regulated genes** | | | | | |
|---|---|---|---|---|---|
| COG categories | COG & DE | COG but !DE | !COG but DE | !COG&! DE | P-value |
| **CELLULAR PROCESSES AND SIGNALING** | | | | | |
| [D] Cell cycle control, cell division | 2 | 33 | 425 | 3468 | 0.58 |
| [M] Cell wall/ membrane/ envelope biogenesis | 23 | 205 | 404 | 3296 | 0.82 |
| [N] Cell motility | 10 | 101 | 417 | 3400 | 0.64 |
| [O] Post-translational modification | 12 | 129 | 415 | 3372 | 0.41 |
| [T] Signal transduction mechanisms | 18 | 160 | 409 | 3341 | 0.9 |
| [U] Intracellular trafficking and secretion | 11 | 116 | 416 | 3385 | 0.47 |
| [V] Defense mechanisms | 0 | 47 | 427 | 3454 | 0.007 |
| **INFORMATION STORAGE AND PROCESSING** | | | | | |
| [J] Translation | 45 | 139 | 382 | 3362 | 7.11E-08 |
| [K] Transcription | 31 | 272 | 396 | 3229 | 0.77 |
| [L] Replication, recombination and repair | 31 | 174 | 396 | 3327 | 0.04986 |
| **METABOLISM** | | | | | |
| [C] Energy production and conversion | 27 | 260 | 400 | 3241 | 0.49 |
| [E] Amino acid transport and metabolism | 50 | 312 | 377 | 3189 | 0.06 |
| [F] Nucleotide transport and metabolism | 14 | 84 | 413 | 3417 | 0.25 |
| [G] Carbohydrate transport and metabolism | 19 | 350 | 408 | 3151 | 9.697X10 -5 |

| | | | | | |
|---|---|---|---|---|---|
| [H] Coenzyme transport and metabolism | 13 | 143 | 414 | 3358 | 0.7 |
| [I] Lipid transport and metabolism | 8 | 92 | 419 | 3409 | 0.41 |
| [P] Inorganic ion transport and metabolism | 39 | 176 | 388 | 3325 | 0.000981 |
| [Q] Secondary metabolites biosynthesis | 11 | 53 | 416 | 3448 | 0.1 |
| **POORLY CHARACTERIZED** | | | | | |
| [R] General function prediction only | 38 | 363 | 389 | 3138 | 0.39 |
| [S] Function unknown | 25 | 292 | 402 | 3209 | 0.08 |
