## Supplemental Table 6 for "Laboratory Evolution Experiments Help Identify a Predominant Region of Constitutive Stable DNA Replication Initiation"

| **Strain** | **Genotype** | **Source** |
|---|---|---|
| *GJ13519* | MG1655 Δ(argF-lac)U169 | LBG, CDFD |
| **<u>Derivatives of GJ13519</u>** | | |
| *ΔrnhA* | Δ*rnhA*::FRT | LBG, CDFD |
| *ΔdnaA/dnaA*$^+$ | *ΔdnaA*::FRT/pHYD2388 | LBG, CDFD |
| *ΔrnhA-dnaA/dnaA*$^+$ | *ΔdnaA*::FRT *ΔrnhA*::FRT/pHYD2388 | LBG, CDFD |
| *ΔrnhA-ΔdnaA* | ΔdnaA::FRT ΔrnhA::FRT | This study |
| *GJ13519/dnaA*$^+$ | GJ13519/pHYD2388 | This study |
| *GJ13519ΔhotH/dnaA*$^+$ | GJ13519 Δ/pHYD2388 | This study |
| *ΔrnhAΔdnaAΔhotH/dnaA*$^+$ | *ΔdnaA*::FRT *ΔrnhA*::FRT Δ4555284: 45660615(uxuR-yjiN) /pHYD2388 | This study |
| *ΔrnhA-ΔdnaA-ΔrnhA-ΔdnaAΔhotH* | *ΔdnaA*::FRT *ΔrnhA*::FRT ΔΔ4555284: 45660615(uxuR-yjiN) | This study |
| **Plasmid** | **Description** | **Source** |
| *pUA139* | Low copy plasmid with fast folding GFP mut2 | SAFS lab |
| *pUA139::Wt rrnD IGR* | pUA139 vector carrying 598bp *rrsD-yrdA* intergenic region sequence | This study |
| *pUA139::Mut rrnD IGR* | pUA139 vector carrying 598bp rrsD-yrdA mutant intergenic [G-A(3,429,052), +A(3,429,054)] sequence | This study |
| *pHYD2388* | *pMU575* derivative carrying *S.enterica dnaA*+ | LBG, CDFD |
| **Primer Description** | **Sequence 5' -----> 3'** | **Source** |
| *HotH*KO *pKD13* F | GTTGACGATATTTATTTTGATGGCTATCTGTTTGA Tgtgtaggctggagctgcttcg | This study |
| *HotH*KO *pKD13* R | TTCGCTGGCTGGAGAGCGAGCATCCACTGAAAG CCAattccggggatccgtcgacc | This study |
| *rrsD -yrdA* IGR F | ATTACTCGAGTCGTCAGCGAAACAGCAA | This study |
| *rrsD-yrdA* IGR R | TAATAGATCTGTATGGGCGTAAAACATC | This study |
