## Supplementary figures and images for "Laboratory Evolution Experiments Help Identify a Predominant Region of Constitutive Stable DNA Replication Initiation"

### Supplemental Figure 1

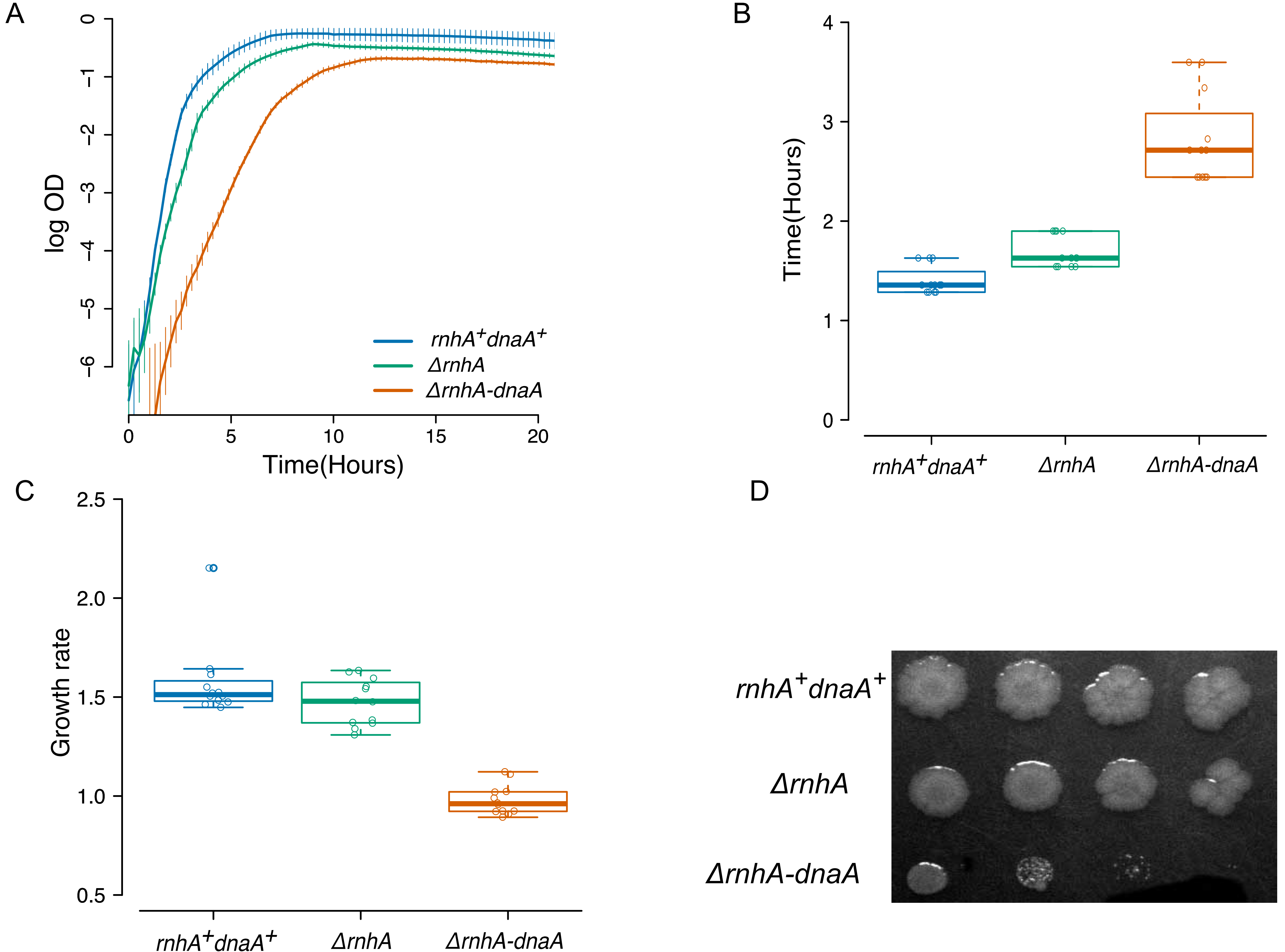

### Supplemental Figure 2

A

B

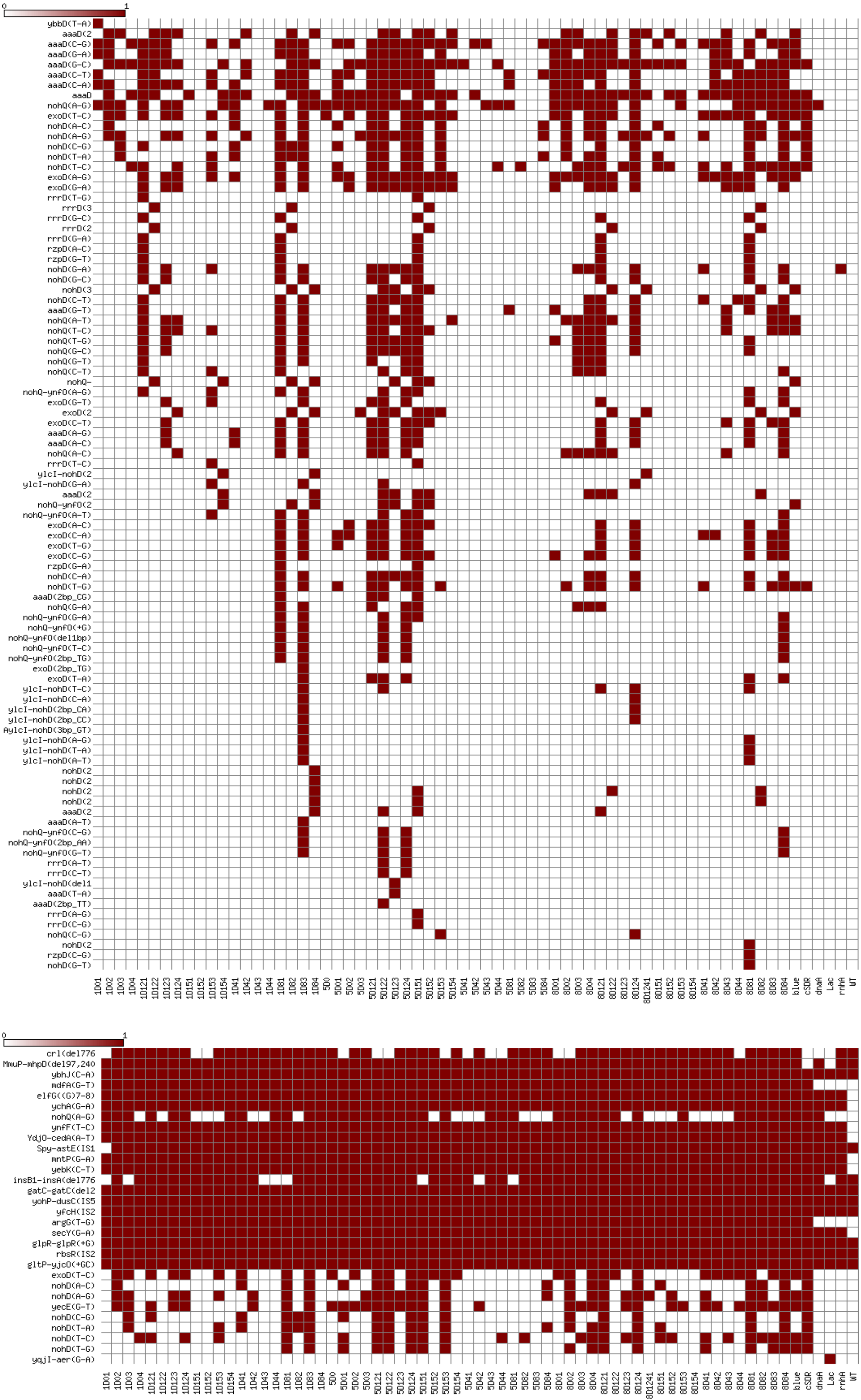

### Supplemental Figure 3

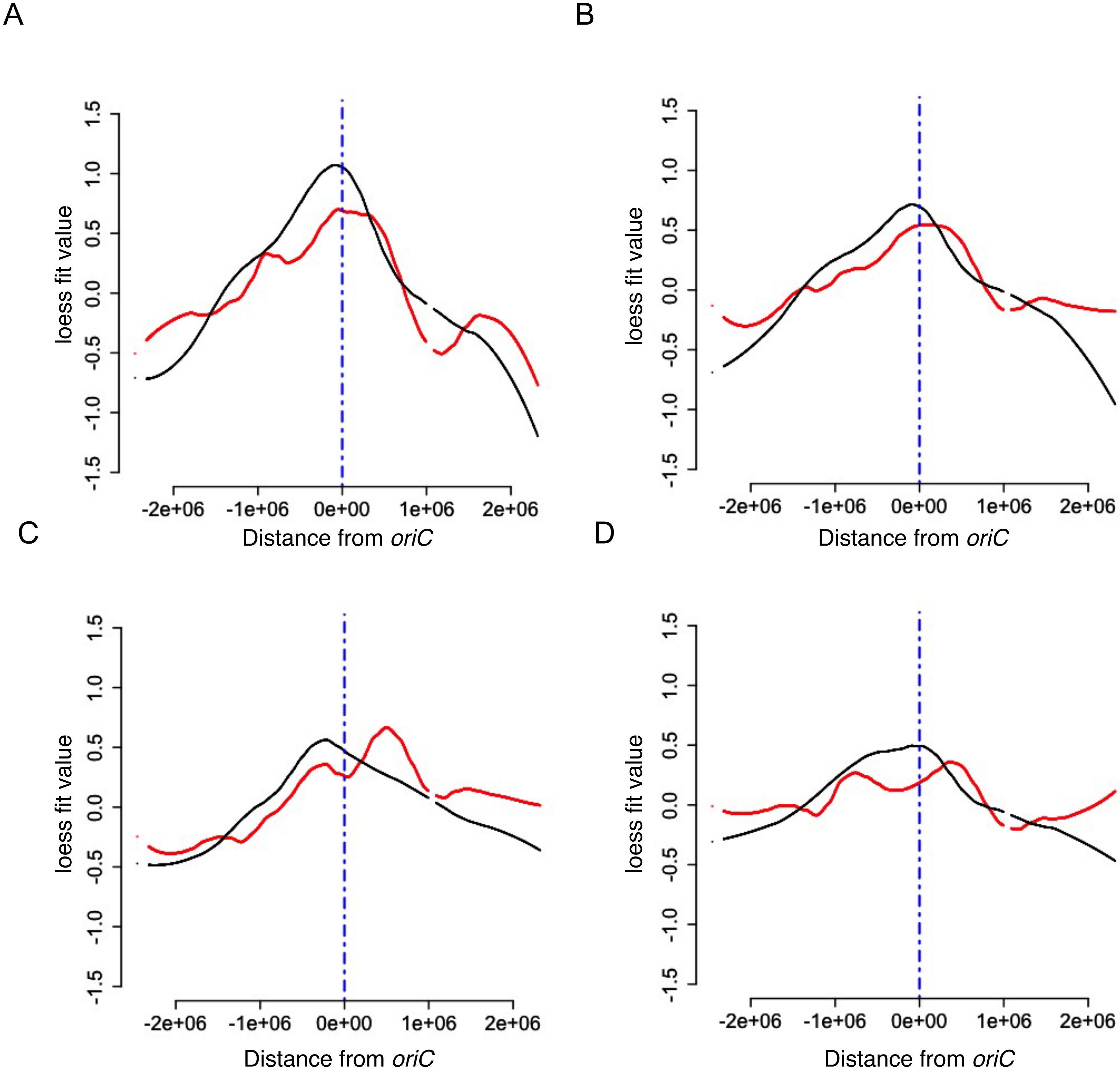

### Supplemental Figure 4

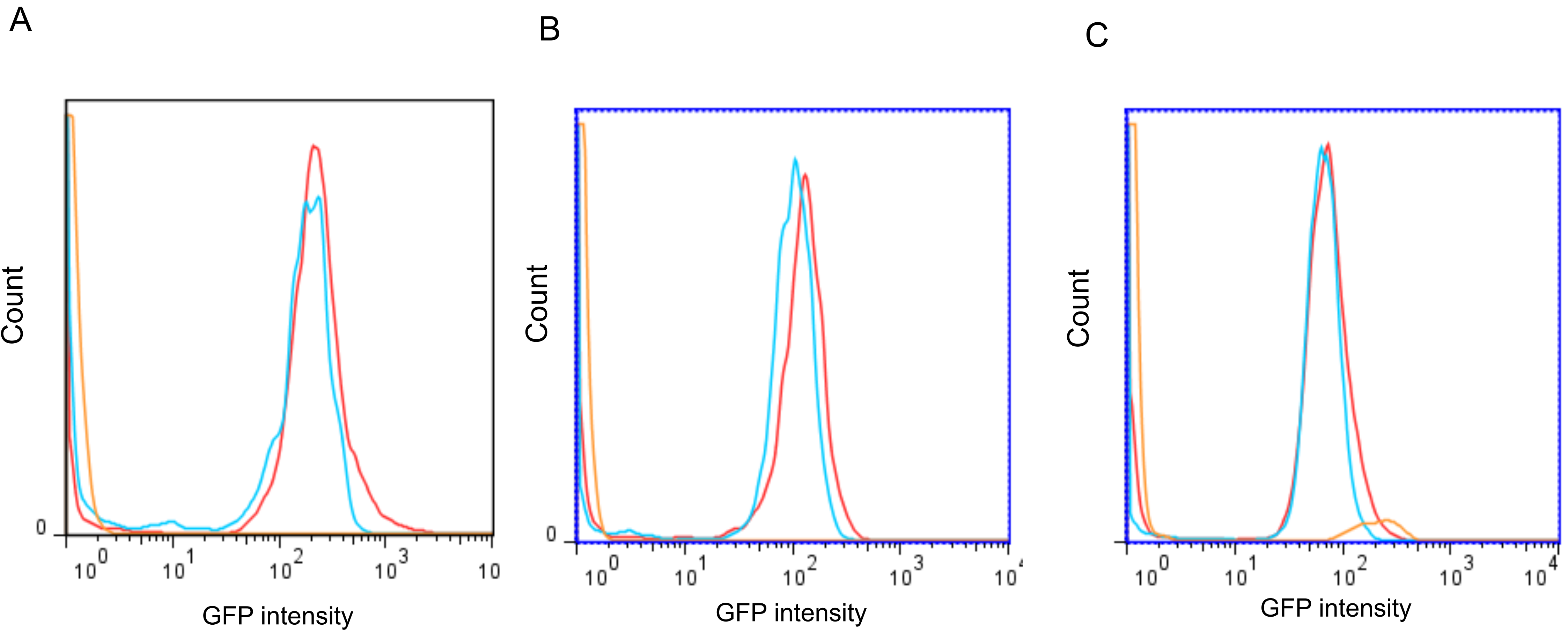
